# Mesoscale liquid model of chromatin recapitulates nuclear order of eukaryotes

**DOI:** 10.1101/634980

**Authors:** Rabia Laghmach, Michele Di Pierro, Davit A Potoyan

## Abstract

The nuclear envelope segregates the genome of Eukaryota from the cytoplasm. Within the nucleus, chromatin is further compartmentalized into architectures that change throughout the lifetime of the cell. Epigenetic patterns along the chromatin polymer strongly correlate with chromatin compartmentalization and, accordingly, also change during the cell life cycle and at differentiation. Recently, it has been suggested that sub-nuclear chromatin compartmentalization might result from a process of liquid-liquid phase separation orchestrated by the epigenetic marking and operated by proteins that bind to chromatin. Here, we translate these observations into a diffuse interface model of chromatin, which we named MEsoscale Liquid mOdel of Nucleus (MELON). Using this streamlined continuum model of the genome, we study the large-scale rearrangements of chromatin that happen at different stages of the growth and senescence of the cell, and during nuclear inversion events. Particularly, we investigate the role of droplet diffusion, fluctuations, and heterochromatin-lamina interactions during nuclear remodeling. Our results indicate that the physical process of liquid-liquid phase separation, together with surface effects is sufficient to recapitulate much of the large-scale morphology and dynamics of chromatin along the life cycle of cells.

**SIGNIFICANCE STATEMENT:** Eukaryotic chromatin occupies a few micrometers of nuclear space while remaining dynamic and accessible for gene regulation. The physical state of nuclear chromatin is shaped by the juxtaposition of complex, out of equilibrium processes on one hand and intrinsic polymeric aspect of the genome on the other. Recent experiments have revealed a remarkable ability of disordered nuclear proteins to drive liquid-liquid phase separation of chromatin domains. We have built a mesoscale liquid model of nuclear chromatin which allows dissecting the contribution of liquid behavior of chromatin to nuclear order of eukaryotes. Our results show that liquid-liquid phase separation, together with surface effects is sufficient for recapitulating large-scale morphology and dynamics of chromatin at many stages of the nuclear cycle.

## INTRODUCTION

Functional compartmentalization is a ubiquitous hallmark of life; by segregating bio-molecules and their interactions cells achieve specialization and improved efficiency of many of their functions (1, 2). In eukaryotic cells, the genetic material is separated from the cytoplasm by the nuclear membrane within the few cubic micrometers of nuclear space. Within the nuclear boundary, we find further compartmentalization which, however, exists in the absence of membranes (3). The structural organization of chromosomes changes with the cell type and phase of life, chromosomal loci have been observed to move across genomic compartments during cell differentiation (4–6). As a consequence, chromatin compartmentalization is believed to play a role in gene regulation, stochastic cell fate determination (7, 8) and the establishment of stable cellular phenotypes (9, 10).

The first evidence of chromatin sub-nuclear organization in interphase was the discovery of the nucleolus in the early nineteenth century, followed by the discovery of regions in the nucleus with distinct optical properties, which were named heterochromatin and euchromatin (11). Heterochromatin appears dense and slow diffusing, containing regions of chromosomes corresponding to mostly silenced genes. Euchromatin, on the other hand, appears less dense and more mobile, composed of mostly active genes (12, 13). The latest electron microscopy tomography experiments have confirmed the DNA density variations between heterochromatic and euchromatic regions without, however, finding any structural difference between the two types of chromatin (14).

Clear evidence of hierarchical compartmentalization in chromatin at multiple scales has emerged through studies employing DNA-DNA proximity ligation assays. First, two genomic compartments were observed (3), named A and B, which were then refined through higher resolution experiments to reveal the existence of even smaller sub-compartments (10). The A and B compartments appear to be enriched in epigenetic markings correlating with transcriptional activation and silencing respectively. Several studies have suggested that chromatin compartmentalization may result from a process of liquid-liquid phase separation orchestrated by the epigenetic markings which collectively act to remodel chromosomal loci (15–19). The epigenetically driven phase segregation events have shown to emerge right at the nucleosome resolution mediated by protein binding as has been shown in the works by Bascom et al (20–22) and Macpherson et al (23).

Chromatin compartmentalization is also consistent with recent experiments that have revealed the remarkable ability of intrinsically disordered proteins to phase separate and form liquid-like protein-rich droplets (24–27). Akin to oil droplets in a well-shaken bottle of vinaigrette, the protein-rich droplets can appear and disappear according to external triggers as well as divide and undergo fusion by forming larger droplets (28–30). Most importantly, in the latest series of experiments, members of Heterochromatin protein 1 family (HP1) known for regulation of heterochromatin content in the nucleus have been shown to drive liquid-liquid phase separation both *in vivo* and *in vitro* (31, 32). Consequently, protein-induced phase separation of chromosomal domains might constitute a direct physical mechanism for regulating genetic processes in space and time in the nucleus (24, 33).

The global architecture of chromosomes appears indeed to be determined by the interplay between chromatin phase separation and motor activity (15–17, 34, 35). Besides chromatin compartmentalization, the other prominent feature of three-dimensional genome architecture, the topologically associated domains (TADs), also appear to arise through phase separation and DNA extrusion (36). Theoretical models based on polymer dynamics have been successfully used to connect one-dimensional epigenetic information to the three-dimensional architecture of the genome. Indeed, it is possible to predict the structural ensembles of human chromosomes with high accuracy relying exclusively on the information extracted from chromatin immunoprecipitation sequencing (ChIP-seq) (16). The same theoretical framework, composed exclusively of polymer connectivity, motor activity, and micro-phase separation, was shown to successfully explain a wide range of experimental observations about the dynamics of chromosomal loci (17). The sub-diffusive behavior of chromatin together with the heterogeneity of the individual diffusing trajectories (37), the viscoelasticity of the nuclear environment (38), and the coherent motion of DNA observed by correlation spectroscopy (39) were all naturally predicted through theoretical and computational modeling.

Here we set out to investigate the specific contribution of liquid-liquid phase separation to genome architecture and separate its effects from those of polymer connectivity and motor activity. To gain insights into the roles of phase separation and surface effects in chromatin compartmentalization, we introduce the MEsoscale Liquid mOdel of Nucleus (MELON), a physical model of nuclear organization which is rooted in the theory of complex fluids. We use diffuse interface finite element simulations to model the evolution of nuclear chromatin compartments under various developmental processes, including growth and inversion/senescence. Our approach draws inspiration and integrates elements of several mesoscopic cellular models proposed previously. These are the two-fluid fluctuating hydrodynamic model of chromatin (40), deterministic phase-field models of multi-cellular domain growth (41, 42) and its mathematical application to rod chromocenter patterning (43), mesoscale polymeric models of chromatin fiber (20–22) and active cellular mechanics models (44–47). We have applied MELON framework to model liquid-liquid phase separation driven reorganization of *Drosophila melanogaster* nucleus under different conditions which are characteristic for different cell phases (48, 49): interphase, active remodeling phases, long-term senescence and nuclear inversion (Fig 1A). Finally, we note that the generic nature of the MELON framework (Fig 1B) along with the minimal physical assumptions that we have built into the model of Drosophila nucleus allows drawing broad inferences which should also hold for other eukaryotic nuclei.

**Figure 1:**
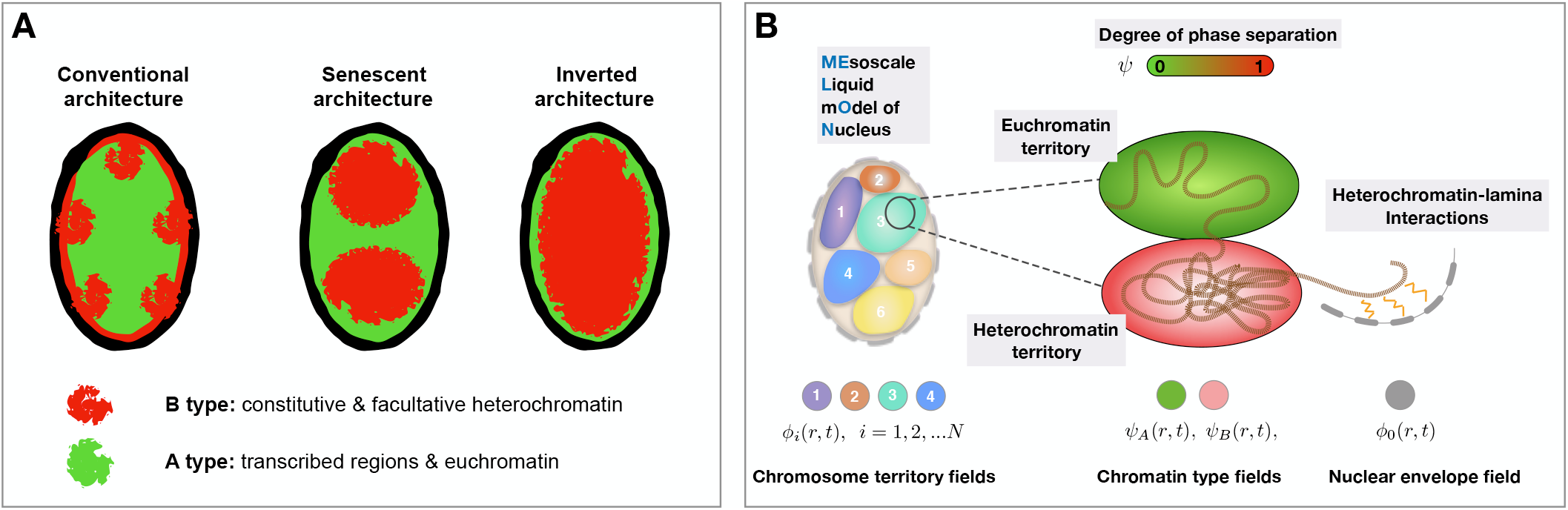
(A) Schematic representation of commonly observed nuclear architectures for *Drosophila* nucleus including conventional, senescent and inverted states observed upon loss of Lamin-heterochromatin tethering (48, 49) (B) Illustration of the MELON framework. The nucleus is resolved by several marginally overlapping liquid-like chromosomal territories, each of which is described by an individual field variable. Within each chromosomal territory, we introduce an additional variable describing (possible) chromatin phase separation into A/B types that form hetero or euchromatin droplets. A separate field variable is introduced for describing the elastic membrane and its relaxation dynamics during the events of nuclear growth or inversion.

## 1 MATERIALS AND METHODS

### The mesoscale liquid model of the nucleus (MELON)

In this section, we present the basic physics and motivating biology behind the MEsoscale Liquid mOdel of Nucleus (MELON). More details about the physical formulation and computational implementation of MELON are in the Supporting Information (SI). The MELON targets modeling (i) long time scale chromatin reorganization dynamics, (ii) liquid-liquid phase separation of chromatin types (ii) fluctuations and other non-equilibrium processed of nuclear remodeling. To this end, we have put together a global free energy functional based on essential physical features of chromatin: phase separation, surface tension, volume constraints and specific interaction of chromatin types. The nuclear chromatin morphology is defined through fluctuating order parameters which resolve (i) Nuclear membrane *ϕ*_0_(*r*, *t*) (ii) Global *i* = 1, …, *N* chromosome territories *ϕ*_*i*_(*r*, *t*) and (iii) Epigenetic states of chromatin *ψ*(*r*, *t*) which smoothly varies from 0 to 1 corresponding to A and B chromatin types respectively (Fig. 1).

The time evolution of all the order parameters is governed by the global free energy functional. The specific forms of the free energy functional terms are motivated either by basic experimental facts about the chromosomal organization (existence of territories and types) or polymeric physics of chromosomes (excluded volume, de-mixing of types). The mesoscale resolution of chromatin naturally accounts for epigenetically driven liquid-liquid phase separation, surface effects of domains as well as diffusion and fluctuations of liquid chromatin droplets within the nucleus.

After all the relevant interactions are accounted, the steady-state nuclear morphology is generated by a stochastic search for the global minimum of the free energy functional in the space of phase-field variables ***φ*** = {*φ*_0_, {*ϕ*_*i*_}_*i*=1,…,*N*_, *ψ*}:

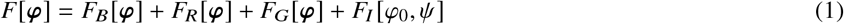

The base free energy term *F*_*B*_ accounts for surface energy contribution of chromosomal interfaces and intra-chromosomal A/B interfaces. For all domains defined by field variables ***φ*** = {*φ*_0_, {*ϕ*_*i*_}_*i*=1,…,*N*_, *ψ*} we assume a simple Landau-Ginzburg form including double-well bulk free energy and surface contributions:

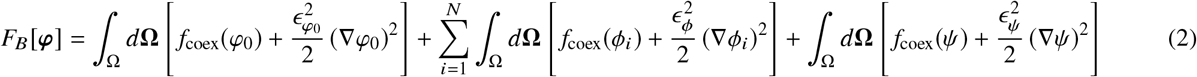

where Ω is the domain of the simulation, and E coefficients are thickness parameters and *f*_*coex*_ is the free energy function which maintains coexistence between two chromosomal phases. The restriction term *F*_*R*_ [***φ***] establishes chromosomal territories in the nucleus by penalizing the spatial overlap between chromosomal domains described by field variables *ϕ*_*i*_.

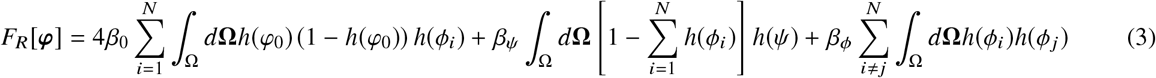

The free energy penalties for excluded volume interactions are introduced via positive volume overlap terms *β*_*ij*_*∫*_Ω_ *d***Ω***h*(*ϕ*_*i*_)*h*(*ϕ*_*j*_) between different domains. The *h*(*φ*_*i*_) functions are standard polynomial forms for approximating volumes of different domains, and can be found in SI. The strength of interaction between different chromatin and nuclear domains are dictated by the energetic prefactors *β*_*ij*_. These prefactors quantify heterochromatin-nuclear envelope soft-excluded volume interactions *β*_*φ*0*ψ*_ = *β*_0_ = *const*, chromosome-chromosome soft-excluded volume interactions *β*_*φiφj*_ = *β*_*φ*_ = *const* and euchromatin-heterochromatin mixing affinity *β*_*ψφi*_ = *β*_*ψ*_ (See SI for numeric values of all the coefficients). The mixing affinity term *β*_*ψ*_ is varied extensively during simulations for investigating the impact of A/B type interactions on nuclear morphology and kinetics of chromatin reorganization during nuclear remodeling processes.

The growth free energy term ***F***_***G***_[***φ***] controls the volume growth/shrinking of chromosomal territories upon nuclear volume changes:

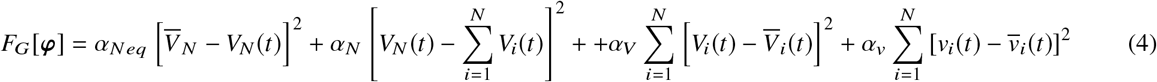

The growth terms are defined via the harmonic restraint 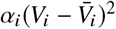 terms which favor stable domain size for the nucleus *V*_*N*_, chromosomal territories 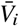 and heterohromal territories 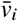. The energetic prefactors α_*i*_ govern the strength of domain localization (See SI for more information). Similar formulation of domain growth terms as volume constraints in free energy has been used by Nonomura in applications of phase-field methods to multi-cellular growth models (41) and recently also by Lee et al (43) for mathematical modeling of chromocenter patterns of rods. Recent experiments and simulations have been shown that nuclear chromatin organization is highly correlated with integrity of chromatin-lamina scaffold (48, 50), which regulates chromatin dynamics during development. To highlight the effect of nuclear shape dynamics on chromatin re-organization it is important to couple the nuclear shape dynamics with chromatin state variables. The *F*_*I*_[*φ*_0_, *ψ*] term accounts for heterochromatin-lamina interactions giving rise to the so-called lamina associating domains (LADs) formed by heterochromatin regions (51). LADs have a significant nuclear presence and are localized near the inner nuclear membranes in most of the mammalian nuclei.

We model this aspect of nuclear architecture by a strong membrane affinity term which keeps heterochromatin preferentially clustered in the vicinity of the membrane region. This preferential interaction of heterochromatin with nuclear lamina is realized via interfacial free energy functional:

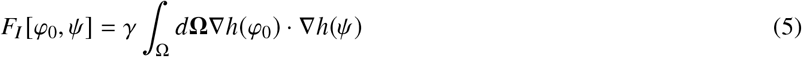

where *γ* is a binding affinity coefficient quantifying how strong heterochromatin is “attracted to” nuclear lamina or nuclear envelope relative to intra-chromosomal interactions. After specifying the full free energy functional of nucleus (Eqs. (1) – (5)) the dynamic equations can be written for each phase-field variables using an Allen-Cahn prescription (52):

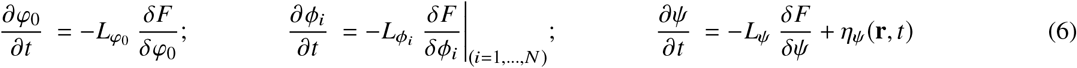

The term *η*_*ψ*_ accounts for fluctuations at the boundaries of euchromatin/heterochromatin islands due to finite size nature of droplets. The fluctuations are modeled as Brownian noise: ⟨*η*_*ψ*_(**r**, *t*)*η*_*ψ*_(**r**′, *t*′)⟩ = *A*_*p*_*δ*(**r−r**′)*δ*(*t−t*) where the amplitude of noise *A*_*p*_ = 2*k*_*B*_*T*_*eff*_*L*_*ψ*_ sets the “effective temperature” *T*_*eff*_ of the nucleus (40) which can be taken as a measure of ATP activity in comparisons with experiment (53, 54). We note that in the MELON framework one can readily introduce active processes and driven fluctuations in chromatin liquid droplets (53, 54). The time scale of chromatin relaxation is set by τ = *L*^−1^. It is well known that in different developmental stages of eukaryotic cells, the dynamics of nuclear processes proceed on vastly different time-scales. Therefore, when modeling nuclear rearrangements in post-embryonic interphase, we set τ = 5 · 10^−3^*h* and when modeling long-term nuclear senescence, we set τ = 5*h* to match the relevant time-scales of attaining different nuclear morphology.

### MELON: a physical simulator of liquid-liquid phase transitions in the nucleus

Here we place the MELON framework in the context of previous efforts in applying continuum diffuse interface methods for modeling cellular phenomena. The two-fluid model of chromatin proposed by Bruinsma et al (40) was the first work suggesting the broad usefulness for field-theoretic models for incorporating thermal fluctuations, active processes and chromatin hydrodynamics under one framework. In recent years, field theoretical methods and the phase field methods in particular have emerged as powerful technique for modeling cellular phenomena from protein coacervation (55) to membrane deformation (56–58) and cell division (41, 42). An important class of models originally developed for modeling multi-cellular growth based on phase-field approach have introduced important numerical techniques for modeling domain growth processes under volume constraints. The domain growth free energy functional pioneered by Nonomura et al (41, 42) in particular has inspired many applications. This functional takes into account different forces such as cell adhesion, volume constraints and excluded volumes interaction. A mathematical application of multi-cellular growth model (41) to single-cell objects was presented in the work of Lee et al (43), where the free energy functional of Nonomura (41) was used for generating chromocenter patterns of rod cells and studying patterns with respect of domain volume adjustment. In this model, the cell shape and size nucleus variation are treated via a mathematical translation operation which is uncoupled to any of internal field variables. Furthermore, mathematical switching functions of phase variables have been utilized to change field variables halfway of pattern rod chromocenter generation. This approach comes with a cost of operating a large number of auxiliary parameters which are manually adjusted for generating each target pattern.

The objective for developing MELON is to have a stochastic physical simulator which models passive fluctuations, liquid-liquid phase separation and lamina-anchoring closely mimicking soft polymeric of entities. Compared to mathematical and deterministic applications of diffuse interphase or related continuum models, there are very few parameters, most of which are fixed and mirror coarse-grained polymeric objects in the nucleus. This is achieved by modifying the standard domain growth free energy functional such as found in Nonomura et al., namely: (i) Membrane envelope is explicitly resolved with a new variable and dynamics is propagated by coupling to constant time-scale viscous relaxation. (ii) New coupling terms between chromosomal domains of the nucleus has been introduced. This later couple all the fields together and with membrane thereby act as dense polymeric material. (iii) Thermal noise which couples A/B type fluctuations in heterochromatin and euchromatin territories.

## 2 RESULTS

### Impact of chromosome territorial affinity and heterochromatin-lamina interactions on euchromatin/heterochromatin phase separation

To illustrate the consequences of liquid behavior and phase separation of A/B chromatin types in the nucleus, we apply MELON to model *Drosophila* nucleus in its various developmental stages (Fig 1A). Recent experiments suggest that liquid-liquid phase separation of chromatin domains in *Drosophila melanogaster* nuclei is orchestrated by HP1a proteins, which under favorable conditions would form liquid condensates and dissolve heterochromatin regions within liquid droplets (31, 32, 59). Polymer models have shown the importance of A/B phase separation (16, 17) as one of the main driving forces behind chromatin organization in mammals. Heterochromatin-lamina interactions (50, 60, 61) are instead thought to be responsible for the switching between conventional and inverted nuclear morphology.

Using our diffuse interface model of the nucleus, we investigate the role of various terms in the global free energy functional in generating steady-state nuclear morphology liquid-liquid phase separation dynamics. We set the average heterochromatin content of the nucleus at 25% corresponding to the post-embryonic stage of *Drosophila* nucleus (62). Later on, we will vary this content when investigating the impact of heterochromatin content on the dynamics of phase separation and the resulting nuclear morphology.

To describe chromatin dynamics in the nucleus quantitatively, we evaluate the temporal evolution of the summed volumes of individual chromosomes *V*(*t*) and heterochromatin droplets from all chromosomes *v*(*t*). The simulations that generate starting nuclear morphologies (Fig. 2A, Supplementary movies 1 and 2) show that volumes *V*(*t*) and *v*(*t*) evolve until nucleus is filled with liquid state chromatin at which point a steady state is reached where compartments acquire well-defined volumes. After coexistence between different liquid chromatin compartments is reached, we investigate how variations of interaction strengths defined in the the previous section alter the coexistence state of the nucleus.

**Figure 2:**
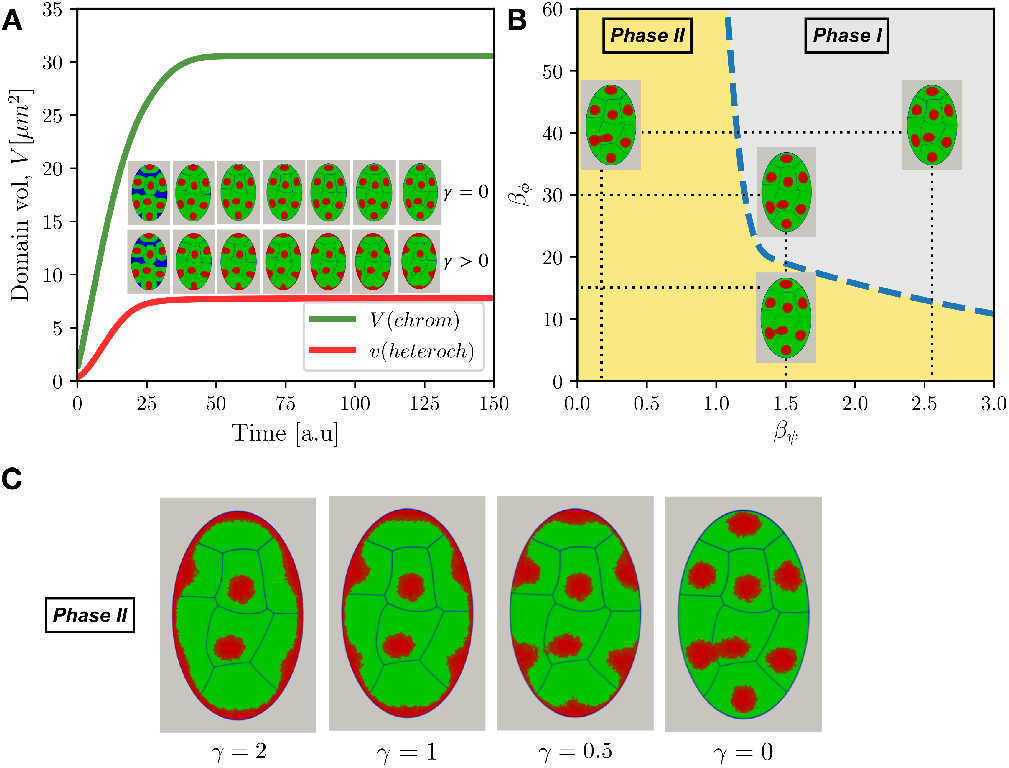
(A) Evolution of chromosomal and heterochromatin volume in the idealized nucleus with and without heterochromatin-lamina interactions (γ > 0 and γ = 0). The snapshots show the nuclear morphology at different times (every 24 time-steps) during the generation stage of the nucleus with and without heterochromatin-lamina interactions. The nuclear compartment is colored in blue, chromosome territory in green and heterochromatin compartment in red. (B) Phase diagram of nuclear morphology showing the impact of various constraints/affinities on liquid-liquid phase separation of A/B domains in the formation of heterochromatin/euchromatin territories. Phases are defined in terms of connectivity between heterochromatin droplets (continuous vs discontinuous or piece-wise continuous variation of *ψ* variable, see also SI). Phase I corresponding to fully disconnected and mixed state phase II corresponding to strongly connected and de-mixed or partially de-mixed states (C) Impact of lamina-heterochromatin anchoring affinity variation on the emergent nuclear morphology.

We first investigate the impact of chromosome type mixing affinity *β*_*ϕ*_ and *β*_*ψ*_ which the later governs the interaction range between chromatin compartments. We see that, for a given chromosomal interaction range *β*_*ϕ*_, a strong affinity leads to mixed states for heterochromatin droplets while weaker mixing affinity leads to fusion of heterochromatin droplets (Fig. 2B).

Variation of nuclear membrane lamina-heterochromatin interaction shows that nuclear morphology is significantly affected by the changes in lamina anchoring strength (Fig. 2C). This is unsurprising given the large surface area of the nuclear envelope. Indeed when we set a non-zero affinity *γ* > 0 for the lamina interactions, the heterochromatin shows pronounced localization near the nuclear envelope. In the presence of dominant lamina-heterochromatin interactions, the impact of mixing affinity is now seen in different thickness of the heterochromatin belt formed around the nuclear envelope. This somewhat subtle difference is due to the competition between the lamina binding energy (negative term in the global *F* [*φ*]) and territorial interaction energy (positive term in the global *F* [*φ*]). Thus, consistent with optical microscopy observations (62, 63) and polymer simulations (50, 60), the lamina binding affinity is indeed one of the dominant physical interaction that shapes conventional nuclear architecture.

Loss of affinity has been shown to lead to an inversion of nuclear architecture, which we show is a naturally emerging behavior of our model of the nucleus (63, 64). In the subsequent sections, we show that the inversion behavior along with its characteristic kinetics is naturally explained by liquid-like behavior of chromatin.

### Impact of diffusion, fluctuations and heterochromatin content on formation of liquid-chromatin compartments

In this section, we study the roles of heterochromatin droplet diffusion, fluctuations and nuclear heterochromatin fraction on the kinetics of phase separation in the idealized nucleus during interphase and senescent phases. The setup of the first set of simulation is mimicking the rapid liquid-like nucleo-protein droplet fusion events during interphase the dynamics of which have been observed and quantified in multiple recent experiments (31, 32, 65). It is acknowledged that within *In vivo* nucleoplasm environment, heterochromatin droplets display shape and size fluctuations as a result of thermal fluctuations and active, ATP-driven motor activity (66). In the present scheme of the MELON framework, we model fluctuations by a thermal noise term which accounts for the fluctuations of finite-sized droplets in an effective manner. Introduction of fluctuation term, however, has allowed us to quantify how “effective temperature” in the nucleus impacts the kinetics of phase separation. To this end, we have carried out simulations with fixed nuclear volume while varying fluctuation amplitude values.

We find that passive fluctuations lead to enhancement of liquid droplet contact and fusion events, which in turn enhance the kinetics of phase separation in the nucleus (Fig. 3A,B). To be more quantitative, we have evaluated the droplet fusion time *τ*_*c*_ for different fluctuation amplitude *A*_*p*_ (Fig. 3A). Thus, the fluctuation amplitude of heterochromatin/euchromatin interfaces effectively increases the capture radius or diffusion coefficient of the liquid droplets (Fig. 3A,B). Finally, the comparison with experiments probing *in vivo* nucleoplasm (65) shows that liquid state model of chromatin in MELON framework accurately capture the nuclear coalescence profiles as well as surface profiles and sequence of steps on time-lapse nuclear domain coalescence events (Fig. 3B,C).

**Figure 3:**
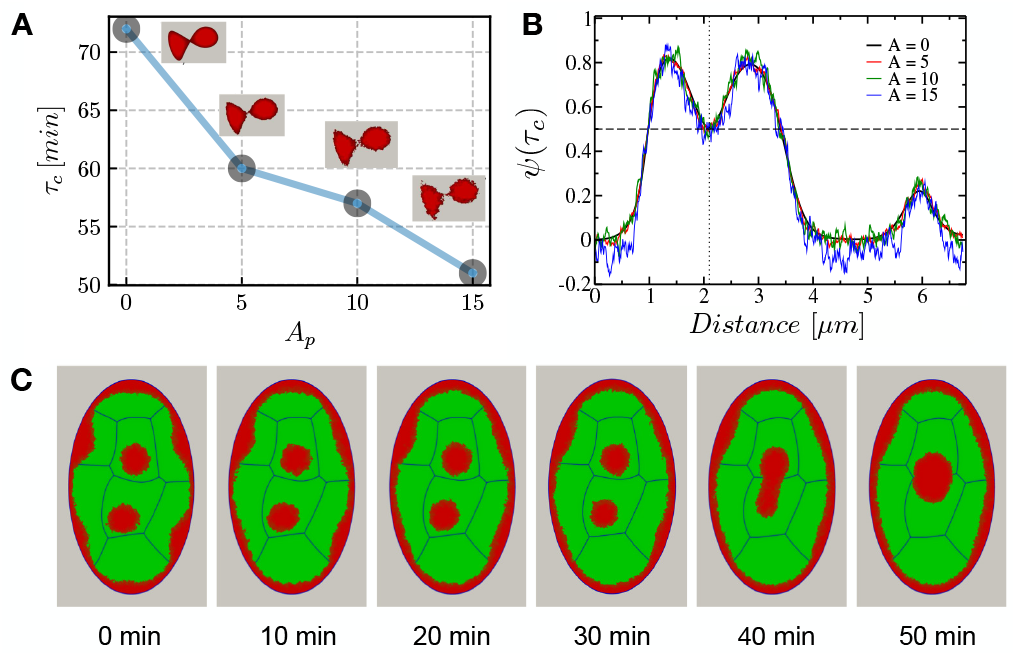
Impact of euchromatin/heterochromatin boundary fluctuations on the heterochromatin droplet fusion dynamics. Shown are (A) Profile of droplet fusion time *τ*_*c*_ as a function of droplet fluctuation amplitude *A*_*p*_ (B) The fluctuation profile along the heterochromatin order parameter (C) Representative snapshot of droplet fusion dynamics for *A_p_* = 10 amplitude and post-embryonic interphase cycle for which chromatin relaxation timescale is set: *τ* = *L*^−1^ = 5 · 10^−3^ *h*

Next, we fix the fluctuation amplitude of droplets at a fixed moderate value and examine a process of large scale nuclear reorganization which is initiated by severing the lamina and heterochromatin. These types of simulations serve as a point of comparison with nuclear volume remodeling simulations of senescence and inversion reported in the next section. Starting from a conventional nuclear morphology with 25% of heterochromatin fraction content, we monitor droplet fusion leading to partial phase separation of heterochromatin. Consistent with the microscopy experiments (62–64), the conventional morphology of the nucleus, in response to the disruption of heterochromatin-lamina anchoring evolves towards morphology with fewer heterochromatic centers which are localized in the center of the nucleus (Fig. 4A). During the reorganization stage, the adjacent heterochromatin droplets fuse, thereby reducing the number of chromocenters in the nucleus (Fig. 4A)). To predict the impact of heterochromatin mobility and its fraction in the nucleus on phase separation kinetics, we performed simulations with different values of heterochromatin fractions and different diffusion coefficients. To do this, we assume that during the nuclear reorganization, the heterochromatin continues growing to reach the prescribed fraction. Indeed, changing the heterochromatin fraction in conventional nuclear architecture to a newly prescribed fraction in inverted architecture should not impact the displacement and fusion of heterochromatin droplets strongly because the growth kinetics of heterochromatin is very fast relative to heterochromatin mobility. Figures 4(B,C) represent the inverted nuclear architecture obtained for 30% and 45% of heterochromatin fraction content in the nucleus. As one would expect, the higher heterochromatin content leads to faster phase separation associated with a decrease in the clustered heterochromatin number.

**Figure 4:**
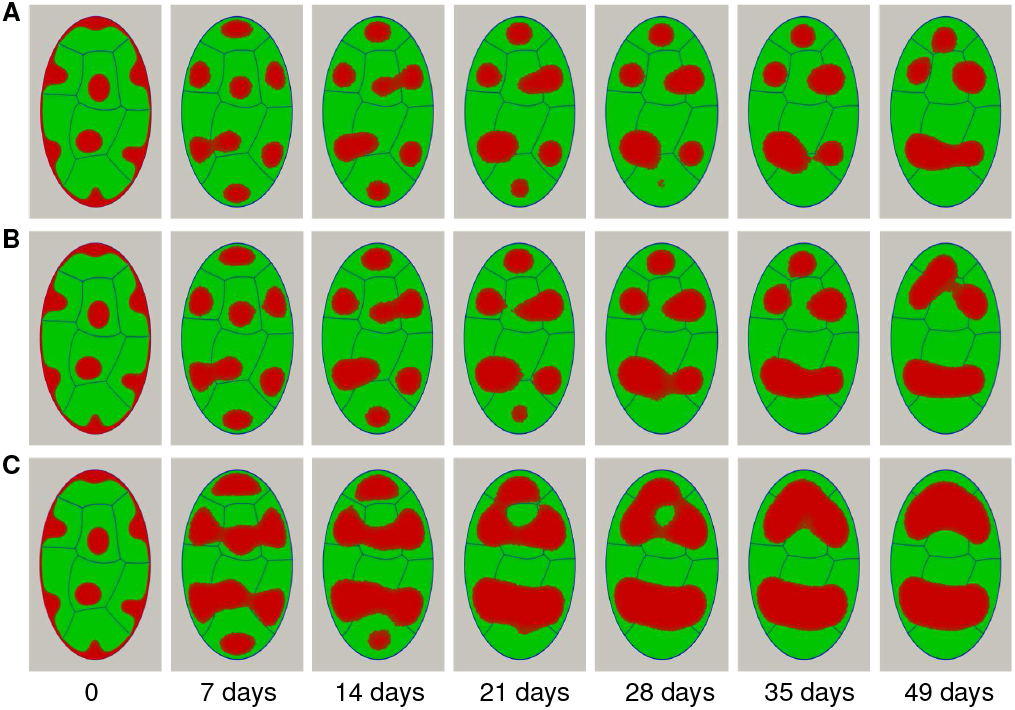
Impact of heterochromatin volume fraction on the nuclear compartment reorganization with fixed nuclear size. Shown are nuclear morphologies generated with different heterochromatin contents: (A) ρ = 25% (B) 30% and (C) ρ = 45%. Simulations are initiated by terminating Lamina-heterochromatin interaction and the time scale of chromatin relaxation is set for modeling nuclear senescence and inversion *τ* = *L*^−1^ = 5*h*. *A*_*p*_ = 5 of mammalian nuclei (62)

The displacement of the heterochromatin droplets within the nucleus is controlled by the diffusion which the velocity is related to the thickness of euchromatin-heterochromatin interfacial region. The simulation results performed for a given heterochromatin fraction (30%) and a fixed nuclear volume showed that increasing the diffusion coefficient accelerate the fusion of heterochromatin droplets to form two clusters at steady-state of nuclear morphology for a higher value (Fig. S3). Additionally, we find there to be a critical threshold exceeding of which leads to fully phase-separated morphology given reasonable diffusion constants and absence of any desegregating lamina-heterochromatin interactions.

### Interplay between chromatin phase separation and nuclear volume remodeling accompanying senescence

The cell nucleus is subject to continuous remodeling activity changing volume, shape or internal organization in response to a variety of external or internal signals. For instance, during the interphase eukaryotic nuclei undergo steady expansion until mitosis (67). During senescence (68) and embryonic development (62, 63, 63), however, the nucleus can lose lamina-heterochromatin affinity and undergo large-scale redistribution of chromatin where heterochromatin moves away from the periphery to a region closer to the center of the nucleus (Fig. 1). This large scale chromatin remodeling events are often accompanied by expanding and shrinking of nuclear volume (63, 63). Given the frequency in which nuclear volume/shape change and chromatin reorganization happen together, one may expect there to be robust mechanisms tuned with volume drift and fluctuations.

Phase separation of chromatin domains provides one such robust mechanism for rapid mobilization of a large section of the genome with predictable dependence on volume changes. In this context, it is, therefore, worthwhile to investigate in detail how the kinetics of chromatin phase separation is coupled with the remodeling of nuclear volume.

To this end, we have carried out simulations that mimic processes of nuclear senescence and inversion happening on the time-scale of days and resulting in a dramatic change in nuclear volume and euchromatin/heterochromatin.

Within the current MELON framework, remodeling of the nuclear volume is simulated by uniaxial slow compression/expansion of membrane boundaries, which allows the liquid chromatin compartments to expand/contract. The volume remodeling processes considered are all propagated at a constant rate which is taken to be much slower than any of the intra-nuclear diffusion time scales: 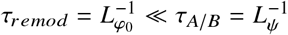. In this work, we did not consider anisotropic deformations and fluctuations of the nuclear envelope (69) which certainly could be an interesting question that merits separate investigation.

The figure 5 shows the impact of nuclear volume changes on chromatin reorganization dynamics in the absence of heterochromatin-lamina interactions. We see that in all cases, heterochromatin/euchromatin distribution driven toward completely phase-separated states with heterochromatin accumulating in the vicinity of the nuclear core. However, the kinetics and morphology of this transition are markedly different relative to constant volume case of the previous section. This shows that chromatin diffusion rates and volume remodeling rates can be in a tug of war with each other. Additionally, we see that the chromosome territories can remain well separated during the remodeling of nuclear volume which is suggesting that coexistence of key sub-nuclear compartmentalization of A/B types with optimal affinity can be made robust to volume remodeling.

**Figure 5:**
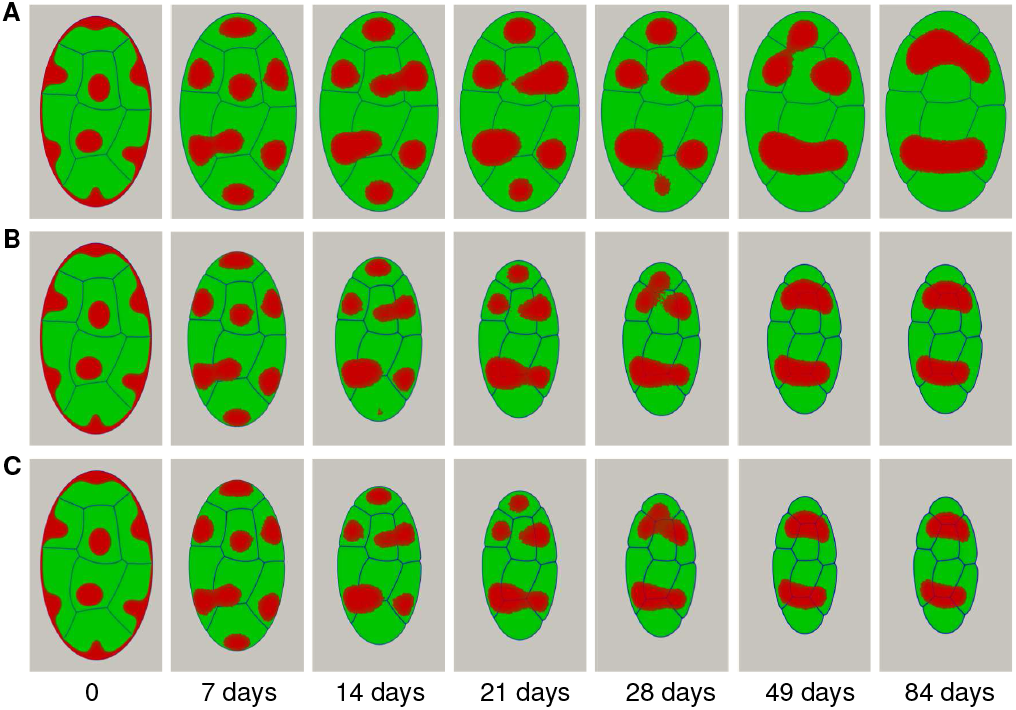
Impact of nuclear volume remodeling on euchromatin/heterochromatin phase separation. Shown are simulations tat remodel nuclear volume by (A) nuclear growth with 10% of volume expansion (B) uniform nuclear compression of oblate nuclear envelope by 50% and (C) Nuclear compression by 60%. Overall heterochromatin fraction content in the nucleus is fixed at 30%.

To quantify the dynamics of heterochromatin/euchromatin phase separation, we have investigated how the number of heterochromatin clusters varies with target nuclear volume while keeping remodeling times fixed. The results show that the value of final volume leads to more rapid phase separation because, in smaller nuclear volumes, the encounter rates of heterochromatin droplets are higher (Fig. 6). Indeed, the distance between the two regions occupied by heterochromatin becomes small when the nuclear volume decreases; hence, the fusion between them becomes more favorable. Since the relaxation time of the nuclear envelope (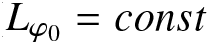) is being kept constant it is also interesting to evaluate the characteristic time of relaxation of heterochromatin domains in order to assess how much the dynamics of internal chromatin motions are affected. We find that the chromosome territories relaxed to their equilibrium volume with a nearly same relaxation time of the nucleus, whereas the heterochromatin domains relaxed faster (Fig 6 and Table S1).

**Figure 6:**
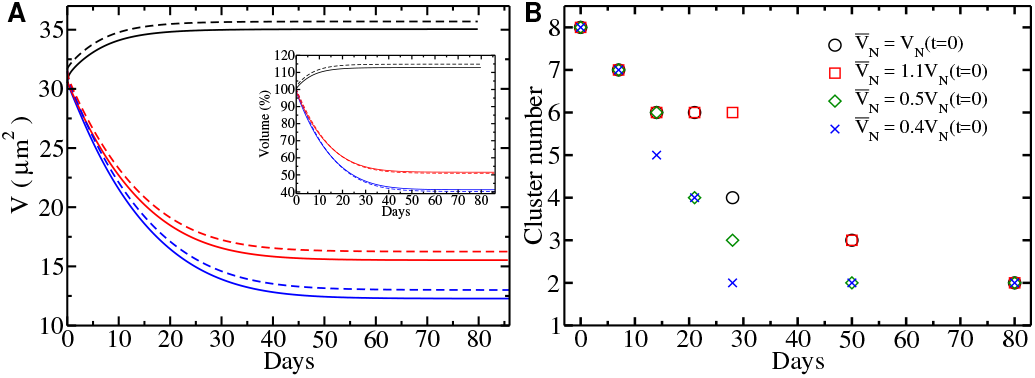
(A) Volume relaxation profiles of chromosomes (solid lines) and nucleus as a whole (dashed lines) as a function of remodeling time. The color code shows the different target of steady-state nuclear volumes corresponding to 10% of volume expansion (black lines), volume compression by 50% (red lines) and volume compression by 60% (blue lines). (B) Temporal evolution of the number of heterochromatin cluster in the cell’s nucleus during the reorganization process with different target steady-state nuclear volumes 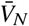. All simulations of the nuclear volume remodeling were initiated by terminating heterochromatin-lamina interaction term to mimic processes of senescence and nuclear inversion.The heterochromatin fraction content in the nucleus is fixed to 30%.

Likewise, the diffusion rates and rate of volume reduction lead to a nearly uniform acceleration of the dynamics of droplet fusion events. We see that nuclear growth works against phase separation and in principle can slow it down or arrest it for some parameter regimes. Based on experimental observations (64), the time required for the complete transformation of the conventional architecture of mammalian nuclei to inverted morphology is predicted to take on the order of ~ 30 days with the corresponding heterochromatin cluster number estimated to be 2-3. The results with embryonic time scale adopted in this work thus agree quantitatively with the experiments (64).

In the case of nuclear inversion, we find that thermodynaic driving force of phase separation ensures robust evolution of chromatin towards steady state nuclear morphologies with respect of heterochromatin content, final volume and diffusion rate variations (Fig. S4, Fig. S5 Fig. S6). This robustness is a generic feature of phase-separations and has been remarked in many related prior studies using phase-field methods such as the ones by Nonomura (41) in the context of multi-cellular growth and Lee et al (43) in the context of rod chromocenters.

Finally, we have explored the conditions under which heterochromatin domains merge into a single cluster which is commonly seen in rod cells upon complete nuclear inversion (Fig. 7). Scanning phase space of nuclear morphology along the heterochromatin density axis (See SI) we have found that specific condition is favoring single cluster formation is having at least 50% heterochromatin content which in the MELON framework would correspond to combined constitutive and facultative heterochromatin content.

**Figure 7:**
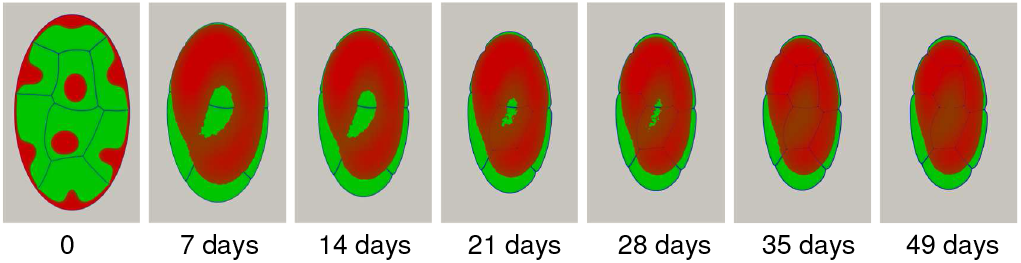
An example of single heterochromatic cluster formation. Heterochromatin fraction in the nucleus is elevated to 60% to account for the existence of high combined facultative and constitutive heterochromatin. We have used ~ 10 times high diffusion coefficient (relative to simulations in Figs 2–6) to accelerate phase separation.

The observations of nuclear inversion has recently attracted considerable attention in studies employing computer simulations and applied mathematics. In particular, we mention here the important work of Falk et al (61) who have used A-B-C hetero-polymeric model of chromatin identifying the balance of type-interaction which leads to inverted architecture. In another work by Lee at al (43), a mathematical application of phase-field methodology based on Nonomura’s multi-cellular growth model (41) was employed to generate inverted nuclear patterns of single rod cells. As opposed to the present work and work by Falk et al (61), however, the inversion was realized via a purely mathematical deterministic protocol of using ad-hoc transnational operators acting on cell envelope and sigmoidal functions adjusting chromatin fractions halfway during protocols. Within the framework of MELON, the nuclear inversion emerges organically as a result of noise-driven dynamical relaxation of chromatin domains upon severing Lamin-heterochromatin anchoring. Furthermore, our simulations indicate the existence of a critical fraction of heterochromatin without which full inversion is hampered.

## CONCLUSION

In recent years there has been a gradual shift away from the paradigm of static regular fiber organization of chromatin to a paradigm of fluid-like, heterogeneous, and dynamic order. This paradigm shift has been further catalyzed by the latest series of experiments which have implicated disordered proteins with known nuclear regulatory functions for driving the liquid-liquid phase separation of chromatin regions in the nucleus. (31, 32, 70, 71). Additionally, several theoretical models (72–75) and computer simulations of chromatin polymer (17, 34, 34, 61, 76) have shown the broad consistency of heterogeneous copolymer melt view of chromatin with the Hi-C and single-molecule data.

Thus, a natural question which emerges from these observations is: how much of large-scale chromatin ordering and dynamics in the nucleus could be explained away just by the fluid-like behavior of chromatin? This question is especially pressing if one wants to create viable models of long-timescale development of eukaryotic nuclei such as in aging, differentiation and disease propagation.

To begin answering these questions, we have constructed MEsoscale Liquid mOdel of Nucleus(MELON); a streamilined computational framework where chromatin is resolved as a fluctuating fluid mixture composed of epigenetically colored components. Using this model, we find that a fluid description of chromatin combined with basic facts about the nuclear architecture, including the existence of chromosomal territories, A/B epigenetic type interactions and Lamina-heterochromatin anchoring leads to life-like nuclear morphologies. Application of the MELON framework to *Drosophila* nucleus at different developmental stages of the nucleus, such as interphase, long-time senescence, and inversion reveals a rich interplay between liquid-liquid phase separation, nucleation, and droplet fluctuations. We would like to emphasize further that the generic nature of the model and the minimal assumptions that we have built into it allows one to draw conclusions which should generally be applicable for a wide variety of eukaryotic nuclei and not just for the *Drosophila* nucleus. Particularly, our study finds that a significant role is played by surface tension and lamina-heterochromatin interactions in determining large-scale chromatin rearrangements. Our study introduces an innovative approach for studying micron-scale chromatin dynamics; we foresee that further development of the method here introduced-the MELON framework-will shed new light on the micro-rheology, the diffusive behavior and the hydrodynamics of nuclear chromatin.

## AUTHOR CONTRIBUTIONS

RL and DAP designed the research. RL carried out all simulations, RL and DAP analyzed the data. RL, MDP and DAP did the literature review and conceptualization. RL, MDP, DAP wrote the article.

## ACKNOWLEDGMENTS

DAP and RL acknowledge useful discussions with Xueyu Song and Jim Evans. DAP is grateful for the financial support from Caldwell foundation of Iowa State. M.D.P.’s research is supported by the Center for Theoretical Biological Physics sponsored by the National Science Foundation (Grants PHY-1427654 and NSF-CHE-1614101) and by the Welch Foundation (Grant C-1792). All the simulations in the paper were carried out using NSF’ XSEDE allocation on Stampede2 machine at the service-provider through allocations: MCB180058 and MCB180071.

## MATERIALS AND METHODS

The MELON framework has been coded in C++ using finite element numerical library MOOSE (77, 78). Nuclear morphology visualizations have been generated via Python-based Paraview (79).

## REFERENCES

1. Alberts, B., 2015. Molecular biology of the cell. Garland Science, Taylor and Francis Group, New York, NY.

2. Phillips, R., 2013. Physical biology of the cell. Garland Science, London New York, NY.

3. Lieberman-Aiden, E., N. L. Van Berkum, L. Williams, M. Imakaev, T. Ragoczy, A. Telling, I. Amit, B. R. Lajoie, P. J. Sabo, M. O. Dorschner, et al., 2009. Comprehensive mapping of long-range interactions reveals folding principles of the human genome. science 326:289–293.

4. Dixon, J. R., I. Jung, S. Selvaraj, Y. Shen, J. E. Antosiewicz-Bourget, A. Y. Lee, Z. Ye, A. Kim, N. Rajagopal, W. Xie, et al., 2015. Chromatin architecture reorganization during stem cell differentiation. Nature 518:331.

5. Chathoth, K. T., and N. R. Zabet, 2019. Chromatin architecture reorganisation during neuronal cell differentiation in Drosophila genome. Genome research gr-246710.

6. Bonev, B., N. M. Cohen, Q. Szabo, L. Fritsch, G. L. Papadopoulos, Y. Lubling, X. Xu, X. Lv, J.-P. Hugnot, A. Tanay, et al., 2017. Multiscale 3D genome rewiring during mouse neural development. Cell 171:557–572.

7. Lin, Y. T., P. G. Hufton, E. J. Lee, and D. A. Potoyan, 2018. A stochastic and dynamical view of pluripotency in mouse embryonic stem cells. PLoS Comp Bio 14:e1006000.

8. Folguera-Blasco, N., R. Pérez-Carrasco, E. Cuyàs, J. A. Menendez, and T. Alarcón, 2019. A multiscale model of epigenetic heterogeneity-driven cell fate decision-making. PLoS Comp Bio 15:e1006592.

9. Meldi, L., and J. H. Brickner, 2011. Compartmentalization of the nucleus. Trends in cell biology 21:701–708.

10. Rao, S. S., M. H. Huntley, N. C. Durand, E. K. Stamenova, I. D. Bochkov, J. T. Robinson, A. L. Sanborn, I. Machol, A. D. Omer, E. S. Lander, et al., 2014. A 3D map of the human genome at kilobase resolution reveals principles of chromatin looping. Cell 159:1665–1680.

11. Heitz, E., 1929. Das Heterochromatin der Moose. Jahrbucher fur Wissenschaftliche Botanik 69:762–818.

12. Imai, R., T. Nozaki, T. Tani, K. Kaizu, K. Hibino, S. Ide, S. Tamura, K. Takahashi, M. Shribak, and K. Maeshima, 2017. Density imaging of heterochromatin in live cells using orientation-independent-DIC microscopy. Mol Biol Cell 28:3349–3359.

13. Grimm, J. B., B. P. English, J. Chen, J. P. Slaughter, Z. Zhang, A. Revyakin, R. Patel, J. J. Macklin, D. Normanno, R. H. Singer, et al., 2015. A general method to improve fluorophores for live-cell and single-molecule microscopy. Nature methods 12:244.

14. Ou, H. D., S. Phan, T. J. Deerinck, A. Thor, M. H. Ellisman, and C. C. O’shea, 2017. ChromEMT: Visualizing 3D chromatin structure and compaction in interphase and mitotic cells. Science 357:370.

15. Di Pierro, M., B. Zhang, E. L. Aiden, P. G. Wolynes, and J. N. Onuchic, 2016. Transferable model for chromosome architecture. Proc. Natl. Acad. Sci. U. S. A. 113:12168–12173.

16. Di Pierro, M., R. R. Cheng, E. L. Aiden, P. G. Wolynes, and J. N. Onuchic, 2017. De novo prediction of human chromosome structures: Epigenetic marking patterns encode genome architecture. Proc. Natl. Acad. Sci. U. S. A. 114:12126–12131.

17. Di Pierro, M., D. A. Potoyan, P. G. Wolynes, and J. N. Onuchic, 2018. Anomalous diffusion, spatial coherence, and viscoelasticity from the energy landscape of human chromosomes. Proc. Natl. Acad. Sci. U. S. A. 115:7753–7758.

18. Barbieri, M., M. Chotalia, J. Fraser, L.-M. Lavitas, J. Dostie, A. Pombo, and M. Nicodemi, 2012. Complexity of chromatin folding is captured by the strings and binders switch model. Proceedings of the National Academy of Sciences 109:16173–16178.

19. Jost, D., P. Carrivain, G. Cavalli, and C. Vaillant, 2014. Modeling epigenome folding: formation and dynamics of topologically associated chromatin domains. Nucleic acids research 42:9553–9561.

20. Bascom, G. D., C. G. Myers, and T. Schlick, 2019. Mesoscale modeling reveals formation of an epigenetically driven HOXC gene hub. Proc. Natl. Acad. Sci. USA 116:4955–4962.

21. Bascom, G. D., and T. Schlick, 2018. Mesoscale Modeling of Chromatin Fibers. In Nuclear Architecture and Dynamics, Elsevier, 123–147.

22. Di Pierro, M., 2019. Inner workings of gene folding. Proc. Natl. Acad. Sci. USA 116:4774–4775.

23. MacPherson, Q., B. Beltran, and A. J. Spakowitz, 2018. Bottom–up modeling of chromatin segregation due to epigenetic modifications. Proc. Natl. Acad. Sci. USA 115:12739–12744.

24. Li, P., S. Banjade, H.-C. Cheng, S. Kim, B. Chen, L. Guo, M. Llaguno, J. V. Hollingsworth, D. S. King, S. F. Banani, et al., 2012. Phase transitions in the assembly of multivalent signalling proteins. Nature 483:336.

25. Brangwynne, C. P., C. R. Eckmann, D. S. Courson, A. Rybarska, C. Hoege, J. Gharakhani, F. Jülicher, and A. A. Hyman, 2009. Germline P granules are liquid droplets that localize by controlled dissolution/condensation. Science 324:1729–1732.

26. Banjade, S., and M. K. Rosen, 2014. Phase transitions of multivalent proteins can promote clustering of membrane receptors. Elife 3:e04123.

27. Brangwynne, C. P., T. J. Mitchison, and A. A. Hyman, 2011. Active liquid-like behavior of nucleoli determines their size and shape in Xenopus laevis oocytes. Proc. Natl. Acad. Sci. USA 108:4334–4339.

28. Brangwynne, C. P., P. Tompa, and R. V. Pappu, 2015. Polymer physics of intracellular phase transitions. Nat Physics 11:899.

29. Uversky, V. N., 2017. Protein intrinsic disorder-based liquid–liquid phase transitions in biological systems: Complex coacervates and membrane-less organelles. Adv. Colloid Interface Sci. 239:97–114.

30. Boeynaems, S., S. Alberti, N. L. Fawzi, T. Mittag, M. Polymenidou, F. Rousseau, J. Schymkowitz, J. Shorter, B. Wolozin, L. Van Den Bosch, et al., 2018. Protein phase separation: a new phase in cell biology. Trends in cell biology 28:420–435.

31. Strom, A. R., A. V. Emelyanov, M. Mir, D. V. Fyodorov, X. Darzacq, and G. H. Karpen, 2017. Phase separation drives heterochromatin domain formation. Nature 547:241.

32. Larson, A. G., D. Elnatan, M. M. Keenen, M. J. Trnka, J. B. Johnston, A. L. Burlingame, D. A. Agard, S. Redding, and G. J. Narlikar, 2017. Liquid droplet formation by HP1α suggests a role for phase separation in heterochromatin. Nature 547:236–240.

33. Shin, Y., and C. P. Brangwynne, 2017. Liquid phase condensation in cell physiology and disease. Science 357:eaaf4382.

34. Liu, L., G. Shi, D. Thirumalai, and C. Hyeon, 2018. Chain organization of human interphase chromosome determines the spatiotemporal dynamics of chromatin loci. PLoS computational biology 14:e1006617.

35. Nuebler, J., G. Fudenberg, M. Imakaev, N. Abdennur, and L. A. Mirny, 2018. Chromatin organization by an interplay of loop extrusion and compartmental segregation. Proceedings of the National Academy of Sciences 115:E6697–E6706.

36. Sanborn, A. L., S. S. P. Rao, S.-C. Huang, N. C. Durand, M. H. Huntley, A. I. Jewett, I. D. Bochkov, D. Chinnappan, A. Cutkosky, J. Li, K. P. Geeting, A. Gnirke, A. Melnikov, D. McKenna, E. K. Stamenova, E. S. Lander, and E. L. Aiden, 2015. Chromatin extrusion explains key features of loop and domain formation in wild-type and engineered genomes. Proc. Natl. Acad. Sci. U. S. A. 112:E6456–65.

37. Bronshtein, I., E. Kepten, I. Kanter, S. Berezin, M. Lindner, A. B. Redwood, S. Mai, S. Gonzalo, R. Foisner, Y. Shav-Tal, and Y. Garini, 2015. Loss of lamin A function increases chromatin dynamics in the nuclear interior. Nat. Commun. 6:8044.

38. Lucas, J. S., Y. Zhang, O. K. Dudko, and C. Murre, 2014. 3D trajectories adopted by coding and regulatory DNA elements: first-passage times for genomic interactions. Cell 158:339–352.

39. Zidovska, A., D. A. Weitz, and T. J. Mitchison, 2013. Micron-scale coherence in interphase chromatin dynamics. Proc. Natl. Acad. Sci. U. S. A. 110:15555–15560.

40. Bruinsma, R., A. Y. Grosberg, Y. Rabin, and A. Zidovska, 2014. Chromatin hydrodynamics. Biophys. J. 106:1871–1881.

41. Nonomura, M., 2012. Study on multicellular systems using a phase field model. PloS one 7:e33501.

42. Akiyama, M., M. Nonomura, A. Tero, and R. Kobayashi, 2018. Numerical study on spindle positioning using phase field method. Physical biology 16:016005.

43. Lee, S. S., S. Tashiro, A. Awazu, and R. Kobayashi, 2017. A new application of the phase-field method for understanding the mechanisms of nuclear architecture reorganization. Journal of mathematical biology 74:333–354.

44. Ghaffarizadeh, A., R. Heiland, S. H. Friedman, S. M. Mumenthaler, and P. Macklin, 2018. PhysiCell: an open source physics-based cell simulator for 3-D multicellular systems. PLoS Comp Bio 14:e1005991.

45. Deviri, D., D. E. Discher, and S. A. Safran, 2017. Rupture Dynamics and Chromatin Herniation in Deformed Nuclei. Biophys. J. 113:1060–1071.

46. Sen, S., A. J. Engler, and D. E. Discher, 2009. Matrix strains induced by cells: computing how far cells can feel. Cell Mol Bioeng 2:39–48.

47. Knežević, M., H. Jiang, and S. Wang, 2018. Active tuning of synaptic patterns enhances immune discrimination. Phys Rev Lett 121:238101.

48. Ulianov, S. V., S. A. Doronin, E. E. Khrameeva, P. I. Kos, A. V. Luzhin, S. S. Starikov, A. A. Galitsyna, V. V. Nenasheva, A. A. Ilyin, I. M. Flyamer, et al., 2019. Nuclear lamina integrity is required for proper spatial organization of chromatin in Drosophila. Nature communications 10:1176.

49. Jagannathan, M., R. Cummings, and Y. M. Yamashita, 2019. The modular mechanism of chromocenter formation in Drosophila. eLife 8:e43938.

50. Kinney, N. A., I. V. Sharakhov, and A. V. Onufriev, 2018. Chromosome–nuclear envelope attachments affect interphase chromosome territories and entanglement. Epigenetics & chromatin 11:3.

51. van Steensel, B., and A. S. Belmont, 2017. Lamina-Associated Domains: Links with Chromosome Architecture, Heterochromatin, and Gene Repression. Cell 169:780–791.

52. Allen, S. M., and J. Cahn, 1979. microscopic theory for antiphase boundary motion and its application to antiphase domain coarsening. Acta. Metall. 27:1084.

53. Saintillan, D., M. J. Shelley, and A. Zidovska, 2018. Extensile motor activity drives coherent motions in a model of interphase chromatin. Proc. Natl. Acad. Sci. USA 115:11442–11447.

54. Zwicker, D., J. Baumgart, S. Redemann, T. Müller-Reichert, A. A. Hyman, and F. Jülicher, 2018. Positioning of Particles in Active Droplets. Phys. Rev. Lett. 121:158102.

55. McCarty, J., K. T. Delaney, S. P. Danielsen, G. H. Fredrickson, and J.-E. Shea, 2019. Complete Phase Diagram for Liquid–Liquid Phase Separation of Intrinsically Disordered Proteins. The journal of physical chemistry letters 10:1644–1652.

56. Lázaro, G. R., I. Pagonabarraga, and A. Hernández-Machado, 2015. Phase-field theories for mathematical modeling of biological membranes. Chemistry and physics of lipids 185:46–60.

57. Lázaro, G. R., I. Pagonabarraga, and A. Hernández-Machado, 2017. Elastic and dynamic properties of membrane phase-field models. The European Physical Journal E 40:77.

58. Elliott, C. M., and B. Stinner, 2010. Modeling and computation of two phase geometric biomembranes using surface finite elements. Journal of Computational Physics 229:6585–6612.

59. Larson, A. G., and G. J. Narlikar, 2018. The Role of Phase Separation in Heterochromatin Formation, Function, and Regulation. Biochemistry 57:2540–2548.

60. Chiang, M., D. Michieletto, C. A. Brackley, N. Rattanavirotkul, H. Mohammed, D. Marenduzzo, and T. Chandra, 2018. Lamina and Heterochromatin Direct Chromosome Organisation in Senescence and Progeria. bioRxiv 468561.

61. Falk, M., Y. Feodorova, N. Naumova, M. Imakaev, B. R. Lajoie, H. Leonhardt, B. Joffe, J. Dekker, G. Fudenberg, I. Solovei, et al., 2019. Heterochromatin drives compartmentalization of inverted and conventional nuclei. Nature 1.

62. Solovei, I., K. Thanisch, and Y. Feodorova, 2016. How to rule the nucleus: divide et impera. Curr. Opin. Cell Biol. 40:47–59.

63. Solovei, I., A. S. Wang, K. Thanisch, C. S. Schmidt, S. Krebs, M. Zwerger, T. V. Cohen, D. Devys, R. Foisner, L. Peichl, H. Herrmann, H. Blum, D. Engelkamp, C. L. Stewart, H. Leonhardt, and B. Joffe, 2013. LBR and lamin A/C sequentially tether peripheral heterochromatin and inversely regulate differentiation. Cell 152:584–598.

64. Solovei, I., M. Kreysing, C. Lanctôt, S. Kösem, L. Peichl, T. Cremer, J. Guck, and B. Joffe, 2009. Nuclear architecture of rod photoreceptor cells adapts to vision in mammalian evolution. Cell 137:356–368.

65. Caragine, C. M., S. C. Haley, and A. Zidovska, 2018. Surface Fluctuations and Coalescence of Nucleolar Droplets in the Human Cell Nucleus. Phys. Rev. Lett. 121:148101.

66. Chu, F.-Y., S. C. Haley, A. Zidovska, and D. A. Weitz, 2017. On the origin of shape fluctuations of the cell nucleus. Proc. Natl. Acad. Sci. USA 114:10338–10343.

67. Gürsoy, G., Y. Xu, L. A. Kenter, and J. Liang, 2014. Spatial confinement is a major determinant of the folding landscape of human chromosomes. Nucleic Acids Res. 42:8223–30.

68. Lukášová, E., A. Kovařík, and S. Kozubek, 2018. Consequences of lamin B1 and lamin B receptor downregulation in senescence. Cells 7:11.

69. Chu, F.-Y., S. C. Haley, and A. Zidovska, 2017. On the origin of shape fluctuations of the cell nucleus. Proc. Natl. Acad. Sci. USA 114:10338–10343.

70. Feric, M., N. Vaidya, T. S. Harmon, D. M. Mitrea, L. Zhu, T. M. Richardson, R. W. Kriwacki, R. V. Pappu, and C. P. Brangwynne, 2016. Coexisting liquid phases underlie nucleolar subcompartments. Cell 165:1686–1697.

71. Gibbs, E. B., and R. W. Kriwacki, 2018. Linker histones as liquid-like glue for chromatin. Proc. Natl. Acad. Sci. USA 115:11868–11870.

72. Maeshima, K., S. Ide, K. Hibino, and M. Sasai, 2016. Liquid-like behavior of chromatin. Curr Op Gen Dev 37:36–45.

73. Iborra, F. J., 2007. Can visco-elastic phase separation, macromolecular crowding and colloidal physics explain nuclear organisation? Theor. Biol. Med. Model. 4:15.

74. Erdel, F., and K. Rippe, 2018. Formation of Chromatin Subcompartments by Phase Separation. Biophys. J. 114:2262–2270.

75. Polovnikov, K., M. Gherardi, M. Cosentino-Lagomarsino, and M. Tamm, 2018. Fractal folding and medium viscoelasticity contribute jointly to chromosome dynamics. Phys Rev Lett 120:088101.

76. Socol, M., R. Wang, D. Jost, P. Carrivain, C. Vaillant, E. Le Cam, V. Dahirel, C. Normand, K. Bystricky, J.-M. Victor, et al., 2019. Rouse model with transient intramolecular contacts on a timescale of seconds recapitulates folding and fluctuation of yeast chromosomes. Nucleic acids research.

77. Slaughter, A. E., J. W. Peterson, D. R. Gaston, C. J. Permann, D. Andrs, and J. M. Miller, 2015. Continuous integration for concurrent MOOSE framework and application development on GitHub. Journal of Open Research Software 3.

78. Gaston, D. R., C. J. Permann, J. W. Peterson, A. E. Slaughter, D. Andrš, Y. Wang, M. P. Short, D. M. Perez, M. R. Tonks, J. Ortensi, L. Zou, and R. C. Martineau, 2015. Physics-based multiscale coupling for full core nuclear reactor simulation. Annals of Nuclear Energy 84:45–54.

79. Ayachit, U., et al., 2012. The ParaView Guide: A Parallel Visualization Application. Kitware Inc. Technical report, ISBN 978-1-930934-24-5.

