## Supplementary Information for "Mesoscale liquid model of chromatin recapitulates nuclear order of eukaryotes"

### Supporting Information for: "Mesoscale liquid model of chromatin recapitulates intra-nuclear order of eukaryotes."

September 14, 2019

#### 1 Supporting Information

##### 1.1 Description of the MELON model for studying chromatin reorganization at nuclear scales

Here we explain in greater detail the physical set up and numerical implementation of the MELON framework to model the drosophila nucleus. In this framework, the state of the nucleus is resolved using several order parameters describing the local state of chromatin within a physical domain $\Omega$. The nucleus is defined through the phase-field variable $\varphi_0(\mathbf{r}, t)$ which takes 0 inside the nucleus and 1 outside with a smooth variation in-between the regions. The variable $\mathbf{r}$ is the spatial position and $t$ is time. The nuclear envelope (membrane) is represented by a diffuse interfacial layer of the nucleus with a small thickness. The $N$ chromosomal territories inside the nucleus are described by individual phase-field variables $(\{\phi_i(\mathbf{r}, t)\}; i = 1, \cdots, N)$. The chromatin types A and B which form the euchromatin-heterochromatin territories are resolved by defining a field variable $\psi(\mathbf{r}, t)$ which quantifies the epigenetic state of the chromosomal region. Each order parameter takes a constant value in its respective two coexisting phases such that it is 1 within its domain, 0 within all the other domains while varying smoothly between interfacial regions. The nucleus interface position is defined through an iso-contour given by $\Gamma_{\varphi_0}(t) = \{\mathbf{r} \in \Omega, \varphi_0(\mathbf{r}, t) = 1/2\}$. For chromosome and heterochromatin interfaces positions are given by the iso-contours $\Gamma_{\phi_i,\psi}(t) = \{\mathbf{r} \in \Omega, \phi_i(\mathbf{r}, t) = 1/2 \text{ and } \psi(\mathbf{r}, t) = 1/2\}$.

To investigate the micron scale fluid dynamics and pattern formation of epigenetically colored nuclear chromatin, we need to construct the local free energy functional based on different physical features of the system: phase separation, surface tension, volume constraints and specific interactions between different chromatin types. We start by introducing the fundamental Ginzburg-Landau free energy functional which describes the phase separation and the overall shape of multi-phase field variables:

$$F_B[\varphi_0, \{\phi_i\}_{i=1,\ldots,N}, \psi] = \int_\Omega d\mathbf{\Omega} \left[ f_{\text{coex}}(\varphi_0) + \frac{\epsilon_{\varphi_0}^2}{2} (\nabla\varphi_0)^2 \right] + \sum_{i=1}^{N} \int_\Omega d\mathbf{\Omega} \left[ f_{\text{coex}}(\phi_i) + \frac{\epsilon_\phi^2}{2} (\nabla\phi_i)^2 \right] + \int_\Omega d\mathbf{\Omega} \left[ f_{\text{coex}}(\psi) + \frac{\epsilon_\psi^2}{2} (\nabla\psi)^2 \right] \tag{1}$$

where $f_{\text{coex}}$ is the coexisting bulk free energy density and the gradient term represent a contribution to the surface energy of the bulk phases interface. The gradient parameters $\epsilon_{\varphi_0}$, $\epsilon_\phi$ and $\epsilon_\psi$ are controlling the thickness of the interfacial region as defined by the variables $\varphi_0$, $\phi_i$ and $\psi$ respectively, and are taken to be small compared to the length-scale of the nucleus. The bulk free energy density has two minima centered at 0 and 1 each of which corresponds to equilibrium bulk phase. We choose the standard double-well potential to describe the coexisting bulk phases: $f_{\text{coex}}(\varphi_0) = \varphi_0^2(1 - \varphi_0)^2/4$. The form of the coexistence function allows connecting the free energy profile of two bulk phases for each phase variable and makes finite contribution only to the interfacial energy. With this expression, the overall shape of the interfaces is governed only by the interfacial energies which penalize the movement of the interfaces. The coexistence function guarantees the attainment of stable diffuse interfaces separating the coexisting bulk phases. The minimization of the $F_B$ gives the profile of the interface: $1/2 \left[1 - \tanh\left(r/2\sqrt{2}\epsilon_{\cdot}\right)\right]$ with thickness parameter $\epsilon_{\cdot}$ defined above for different interfaces. The motion of interfaces is governed not only by interfacial energy but also by local bulk driving force of the phase transition $\Delta f_{\text{bulk}}(\cdot)$, with $f_{\text{bulk}}(\cdot) = f_{\text{coex}}(\cdot) + \cdots$. We now turn to describing the energetic terms representing the bulk driving forces of the phase separation.

To generate the state of a nucleus which is fully occupied by chromosomal domains it is important to allow chromosomal territories to expand and contract in response to nuclear volume changes in a way a dense polymeric solution would respond. In order to account for the constraints imposed by the nuclear volume we introduce a "growth" term of the global free energy functional $F_G$ as follows:

$$F_G[\varphi_0, \{\phi_i\}_{i=1,...,N}, \psi] = \alpha_{Neq}\left[\overline{V}_N - V_N(t)\right]^2 + \alpha_N\left[V_N(t) - \sum_{i=1}^{N} V_i(t)\right]^2 + \\ +\alpha_V \sum_{i=1}^{N}\left[V_i(t) - \overline{V}_i(t)\right]^2 + \alpha_v \sum_{i=1}^{N}\left[v_i(t) - \overline{v}_i(t)\right]^2 \tag{2}$$

where $V_N$ is the nuclear volume, $\alpha_{Neq}$, $\alpha_N$, $\alpha_V$ and $\alpha_v$ are positive coefficients. The first term accounts for the changes of nuclear volume during nuclear reorganization (growth or inversion) stage during which the nucleus evolves towards a new steady-state volume $\overline{V}_N$. The harmonic terms acting on volumes of chromosomal domains ensures that all the chromosomes fill the nuclear volume entirely $V_N$, with an additional volume constraints ensuring the proper ratio of heterochromatin to chromosome domains $\overline{v}_i$ and $\overline{V}_i$ respectively. Notice that the coefficients $\alpha_{Neq}$, $\alpha_N$, $\alpha_V$ and $\alpha_v$ control the kinetics of growth of different domains to reach the prescribed volumes because their respective driving forces are proportional to these parameters.

The volume of each chromosome denoted by $V_i$ and their corresponding heterochromatin domain volume $v_i$ are defined as the spatial integral of their interface profile described by the variables $\phi_i$ and $\psi$. This usual approximation of the volume will change the minima of $f_{\text{bulk}}$ which corresponds to the equilibrium bulk phases. We thus adopt an interpolation function $h$ to approximate the volumes for different compartments while keeping the positions of minima of $f_{\text{bulk}}$ function at 0 and 1. The most frequently adopted polynomial function for calculations using diffuse interface methods [1] is: $h(\varphi_0) = \varphi_0^3(10 - 15\varphi_0 + 6\varphi_0^2)$. Therefore, we approximate the volumes for nucleus $V_N$, chromosomes $V_i$ and heterochromatin domains $v_i$ by:

$$V_N(t) = \int_\Omega d\mathbf{\Omega}\left[1 - h(\varphi_0)\right]; \quad V_i(t) = \int_\Omega d\mathbf{\Omega} h(\phi_i); \qquad v_i(t) = \int_\Omega d\mathbf{\Omega} h(\phi_i)h(\psi)$$

At this point, the free energy functional sketched above describes only the growth and shrinkage of chromosomal domains. To ensure that changes in volumes of chromosomes are coupled to each other we introduce additional energetic terms which penalize the overlap between chromatin domains during the process of nuclear volume change:

$$F_R[\varphi_0, \{\phi_i\}_{i=1,...,N}, \psi] = 4\beta_0 \sum_{i=1}^{N} \int_\Omega d\mathbf{\Omega} h(\varphi_0)\left(1 - h(\varphi_0)\right) h(\phi_i) + \\ \beta_\psi \int_\Omega d\mathbf{\Omega}\left[1 - \sum_{i=1}^{N} h(\phi_i)\right] h(\psi) + \beta_\phi \sum_{i \neq j}^{N} \int_\Omega d\mathbf{\Omega} h(\phi_i)h(\phi_j) \tag{3}$$

Coefficients $\beta_0$, $\beta_\psi$ and $\beta_\phi$ are all positive constants in the model. The first term represents an energetic belt which surrounds the nucleus by restricting the movement of all chromosome territories within the nuclear envelope. In fact, this energy increases when the chromosome territories are near the nuclear envelope and thus penalize strongly the movement of at the peripheral regions of the nucleus. The second and third terms account for the excluded volume interactions between chromosome-chromosome territories and chromatin-heterochromatin domains. Thus, the chromosome territories cannot overlap and maintain their prescribed volume defined previously in $F_G$. The interaction between the heterochromatin and the lamina located at the nuclear envelope plays a crucial role in the nuclear organization and restructuring through the adhesion/desperation processes happening during the cell development. To account for the affinity of heterochromatin towards nuclear envelope, several formulations of adhesion could be developed as a function of corresponding phase-fields variables $\psi$ and $\varphi_0$. Some of the commonly used formulations for adhesive interaction have used polynomial [2, 3] and exponential [3] forms. The adhesion energy depends on the distance of heterochromatin interface from the nuclear envelope. The form of interaction potential between heterochromatin and the nuclear envelope that we have used is the same one which was also used in many prior works of phase-field models [4, 5]:

$$F_I[\varphi_0, \psi] = \gamma \int_\Omega d\mathbf{\Omega} \nabla h(\varphi_0) \cdot \nabla h(\psi) \tag{4}$$

This form of the potential ensures that the adhesion energy is negative in the region of overlapping of the nuclear envelope and heterochromatin interfacial layer and zero elsewhere. Here the $\gamma$ is a positive coefficient which quantifies the binding affinity of heterochromatin domains to nuclear Lamina.

The total free energy of the system that accounting all effect considered for the nature of intra-nuclear chromatin is expressed then as:

$$F[\varphi_0, \{\phi_i\}_{i=1,...,N}, \psi] = F_B + F_G + F_R + F_I \tag{5}$$

The dynamics of chromatin is described through the temporal evolution of the non-conserved order parameters following the Allen-Cahn kinetic equation [6] which is derived through minimization of the free energy functional [5]:

$$\frac{\partial \varphi_0}{\partial t} = -L_{\varphi_0} \frac{\delta F}{\delta \varphi_0}; \qquad \frac{\partial \phi_i}{\partial t} = -L_{\phi_i} \frac{\delta F}{\delta \phi_i}\bigg|_{(i=1,\dots,N)}; \qquad \frac{\partial \psi}{\partial t} = -L_{\psi} \frac{\delta F}{\delta \psi} + \eta_\psi(\mathbf{r},t) \tag{6}$$

where noise terms are defined via fluctuation-dissipation condition $\langle \eta_\psi(\mathbf{r},t)\,\eta_\psi(\mathbf{r}',t')\rangle = 2k_bTL_\psi\delta(\mathbf{r}-\mathbf{r}')\,\delta(t-t')$ with terms $L_{\varphi_0}$, $L_{\phi_i}$ and $L_\psi$ being the mobility coefficients associated with their respective fields. The term $\eta_\psi$ is added to the evolution equation of the phase-field $\psi$ to account the thermal fluctuation effects of the heterochromatin region at the average time-scale. The amplitude of fluctuation is determined by the fluctuation-dissipation relation, which is related to the mobility coefficient $L_\psi$. By calculating the variational derivative of the total free energy functional $F$ in [6], we get the system of relaxation equations:

$$\begin{aligned}\frac{\partial \varphi_0}{\partial t} = \quad & L_{\varphi_0}\Bigg(\epsilon^2_{\varphi_0}\nabla^2\varphi_0 - f'_{\text{coex}}(\varphi_0) - 2h'(\varphi_0)\Bigg\{2\beta_0\,(1-2h(\varphi_0))\sum_{i=1}^{N} h(\phi_i) - \alpha_0\left[V_N(t) - \sum_{i=1}^{N} V_i(t)\right] \\ & -\frac{\gamma}{2}\nabla^2 h(\psi) + \alpha_4\left[\overline{V}_N - V_N(t)\right]\Bigg\}\Bigg)\end{aligned} \tag{7}$$

$$\begin{aligned}\frac{\partial \phi_i}{\partial t} = \quad & L_{\phi_i}\Bigg(\epsilon^2_{\phi}\nabla^2\phi_i - f'_{\text{coex}}(\phi_i) - 2h'(\phi_i)\Bigg\{\alpha_V\left[V_i(t) - \overline{V}_i(t)\right] - \alpha_0\left[V_N(t) - \sum_{i=1}^{N} V_i(t)\right] \\ & +\alpha_v\left[v_i(t) - \overline{v}_i(t)\right]h(\psi) + 2\beta_0 h(\varphi_0)\,(1-h(\varphi_0)) - \frac{\beta_\psi}{2}h(\psi) + \frac{\beta_\phi}{2}\left[\sum_{i=1}^{N} h(\phi_i) - h(\phi_i)\right]\Bigg\}\Bigg)\end{aligned} \tag{8}$$

$$\frac{\partial \psi}{\partial t} = L_\psi\left(\epsilon^2_\psi\nabla^2\psi - f'_{\text{coex}}(\psi) - h'(\psi)\left\{2\alpha_v\sum_{i=1}^{N}\left[v_i(t) - \overline{v}_i(t)\right]h(\phi_i) + \beta_\psi\left(1 - \sum_{i=1}^{N} h(\phi_i)\right) - \gamma\nabla^2 h(\varphi_0)\right\}\right) + \eta_\psi(\mathbf{r},t) \tag{9}$$

When the nuclear volume is constant $V_N(t) = \overline{V}_N$, the governing equations of chromatin dynamics [Eq 6] are reduced to [Eq 8, 9] which also makes the interfacial profile of nucleus independent of time: $\varphi_0(\mathbf{r},t) = \varphi_0(\mathbf{r}) = \frac{1}{2}\left\{1 - \tanh\left(r/2\sqrt{2}\epsilon_{\varphi_0}\right)\right\}$.

Finally, the equations governing chromatin dynamics are resolved in nondimensional form. For that we introduce the length-scale $l$ and the time-scale $t$ for dimensionlizing the equations. The length scale was set to $1\mu m$ and the mobility parameter was chosen to set the time length as a unit.

#### Numerical simulations and parameter setting:

Let consider the application of MELON framework to model drosophila nucleus with 8 chromosome territories. The governing equations of the chromatin dynamics are highly nonlinear. To solve the coupled non-dimensional equations resulting from minimization of the global free energy functional, we use a fully implicit finite element method combined with the preconditioned Jacobian Free Newton Krylov (JFNK) technique [7]. The development of the MELON framework and equations that are derived from it have been implemented in the MOOSE library [8, 9]: it's a massively parallel finite element open-source coded on C++ language which was builds using several high-performance computational libraries for solving nonlinear partial differential equations: MPI, LibMesh, PETSc. Once all the equations are transformed in the weak form to extract the residual vectors, we compute the solution of the phase field variables by using the different kernels implemented in Moose phase-field module. More details of the numerical implementation of the phase-field module are given in [10, 8].

The simulations were performed on a rectangular domain of dimensions 6 $\mu m$ × 9 $\mu m$. A quadrilateral element with 4 nodes was used for domain meshing with the refinement. The total number of elements used for the fine mesh is 640000. The time step of integration $\Delta t$ is fixed at 0.0025 [a.u.]. An elliptical nucleus is introduced in the center of computational domain in which the chromosomes and heterochromatin domains are randomly generated. The initial conditions to generate the conventional nuclear architecture are provided by a tanh-like function: $1/2\left[1 - \tanh\left(r/2\sqrt{2}\epsilon_.\right)\right]$. The geometries of different subdomains described by the phase-field variables $\varphi_0, \{\phi_i\}_{i=1,\dots,N}, and \psi$ have an ellipical shape where $r$ represents the signed distance function. We use the steady-state of conventional architecture as an initial condition to study the nuclear reorganization process. For interface geometry of the nucleus, two cases have been considered in this work. With fixed nucleus volume: $\varphi_0(r)$ and with a dynamic relaxation of the nuclear interface through $\varphi_0(r,t)$ which remodels nuclear volume as function time until steady-state volume $\overline{V}_N$. When nuclear volume is kept constant, the nucleus interface is simulated by an elliptical shape in which the semi-major axes are fixed to $a = 2.5\,\mu m$ and the semi-minor axis to $b = 4\,\mu m$; thus the nuclear

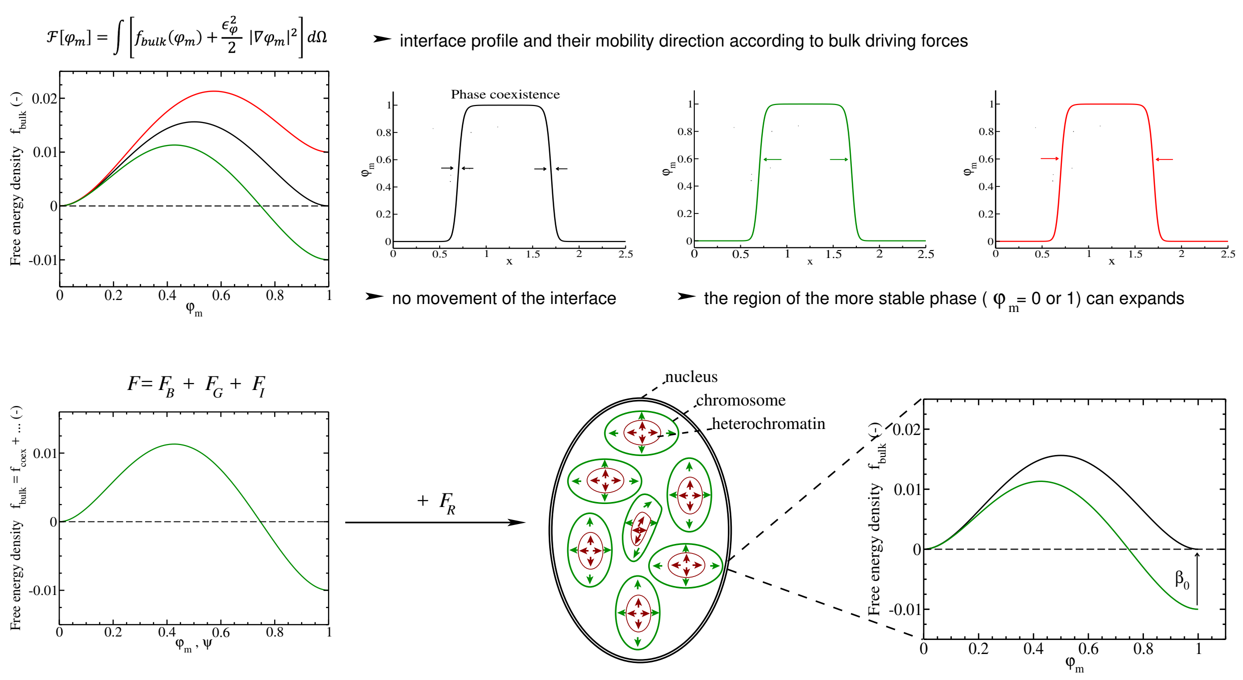


Figure 1: Illustration of MELON free energy terms for intra-nuclear chromatin dynamic. Shown are bulk free energy density as a function of the phase field variable $\phi_m$ and their corresponding interface profile. The relative position of the $f_{bulk}$ indicates the phase transition direction.

volume is: $V_N = \pi ab = 31.416\mu m^2$. For volume remodeling simulations, we compute the temporal evolution of the nuclear volume through h-function as follows: $V_N(t) = \int_\Omega d\mathbf{\Omega}[1 - h(\varphi_0)]$. The mobility coefficient of the nuclear interface is linked to the volume reduction rate in the following way: $L_{\varphi_0} = (V_N(t_0) - \overline{V}_N)/((t_f - t_0) \times V_N(t_0))$, where the coefficient $\alpha_{Neq}$ (fixed at 2 for conventional architecture) controls the nuclear interface velocity . The mobility of chromosomal and heterochromatin compartments are set to be equal $L = L_\varphi = L_\psi$. Since this parameter fixes the inter and intra-chromosomal interface relaxation time: $\tau \propto L^{-1}$. The interface relaxation time $\tau$ is used to set the unit timescale for the kinetics of phase transitions. We note that the value of $\tau$ is naturally expected to be different for different developmental stages of the nucleus. For instance for the post-embryonic interphase of drosophila [11] we set $\tau \sim 0.005\,h$ while for studying long time senescence and nuclear inversion the time scale is calibrated to match different set of experiments [12] and corresponds to $\tau \sim 5\,h$.
The interfacial parameters $\epsilon_\varphi, \epsilon_\psi$ quantify thickness of inter and intra-chromosomal compartments respectively. These parameters can be estimated through the interfacial energies of chromosome territories and euchromatin-heterochromatin compartments. We estimate the interfacial parameters through the diffusion coefficient: $D_\varphi = L\,\epsilon_\varphi^2$ for chromosomes and $D_\psi = L\,\epsilon_\psi^2$ for the heterochromatin regions. By varying the interfacial parameters, we have investigated how the diffusion coefficient impact the nuclear architecture organization and chromatin dynamics. We estimate diffusion coefficients from long-time scale measurements of diffusion of chromosomal loci where clear diffusive behaviour is seen [13, 14]. In this study we have varied the diffusion coefficients wihtin a resonable range that accouts for cellular variablility. The range of values considered is $D_\psi$=[ $0.003\,\mu m^2/h$, $0.004\,\mu m^2/h$, $0.008\,\mu m^2/h$] and $D_\varphi$=[ $0.0015\,\mu m^2/h$, $0.001\,\mu m^2/h$]. The thickness of the diffuse interfacial layer of the nucleus, the heterochromatin and the chromosome compartments are all determined by $\kappa_. = 4\sqrt{2}\epsilon_.\tanh^{-1}(1-2\zeta)$ where $\zeta$ is set to 0.1. The specific values are, $\kappa_{\varphi_0} = 0.44\mu m$ , $\kappa_\phi = 0.44\mu m, 0.538\mu m$ and $\kappa_\psi = 0.761\mu m, 0.88\mu m, 1.243\mu m$. The nondimensionalized parameters of the model such as the intensity of the energetic belt surrounding the nuclear envelope and the excluded volume interaction parameters are simply chosen to be strong enough to contain all the chromosome territories within the nucleus and restrict the overlap. The value of parameters are respectively set to $\beta_0 = 16.67$ and $\beta_\varphi = 40$ for all the simulations. The interaction range of heterochromatin regions has been varied within a range: $\beta_\psi = 0.1 - 0.4$ to investigate its impact on the degree of euchromatin-heterochromatin mixing. For nuclear reorganization simulations, the parameter $\beta_\psi$ was fixed to 0.1. The computed values of the growing domains kinetic parameters are set as $\alpha_0 = 0.16$, and $\alpha_v = \alpha_V = 2$. The interaction parameter between heterochromatin and nuclear lamina is chosen to be the smaller value that allows maintaining the binding between them and is set as $\gamma = 0.009\,\mu m^2$. Euchromatin-heterochromatin interfacial fluctuation have been modeled as thermal noise with an amplitude set to $A = 2k_BTL$. Three values of fluctuation amplitude were considered here A=5, 10 and 15 which corresponding to the different effective temperatures in the nucleus.

Table 1: Fit settings used to extract the characteristic time (or relaxation time). Notice that the time necessary to reach the equilibrium state can be evaluated approximately to $3\tau$

| | Final volume | function used for fit | a($\mu m$) | b | $\tau(h)$ |
|---|---|---|---|---|---|
| Nucleus | | | 35.6931 | 0.11043 | 165.058 |
| Chromosomes | 110% | $a(1 - bexp(-t/\tau))$ | 35.043 | 0.115398 | 166.021 |
| Heterochromatin | | | 10.674 | 0.129893 | 147.994 |
| Nucleus | | | 16.0959 | 0.969034 | 295.307 |
| Chromosomes | 50% | $a(1 + bexp(-t/\tau))$ | 15.3689 | 1.03216 | 296.809 |
| Heterochromatin | | | 4.77723 | 0.9523 | 303.584 |
| Nucleus | | | 12.7916 | 1.47789 | 323.406 |
| Chromosomes | 40% | $a(1 + bexp(-t/\tau))$ | 12.0589 | 1.59465 | 324.675 |
| Heterochromatin | | | 3.77556 | 1.47166 | 331.641 |

#### Data Analysis: Kinetic aspects

The density of the heterochromatin at the i-th chromosome can be expressed as: $\rho_i(t) = v_i(t)/V_i(t)$. Thus, the associated euchromatin density is evaluated as $1 - \rho_i(t)$.
The density of the heterochromatin in cell's nucleus: $\rho(t) = v(t)/V(t)$

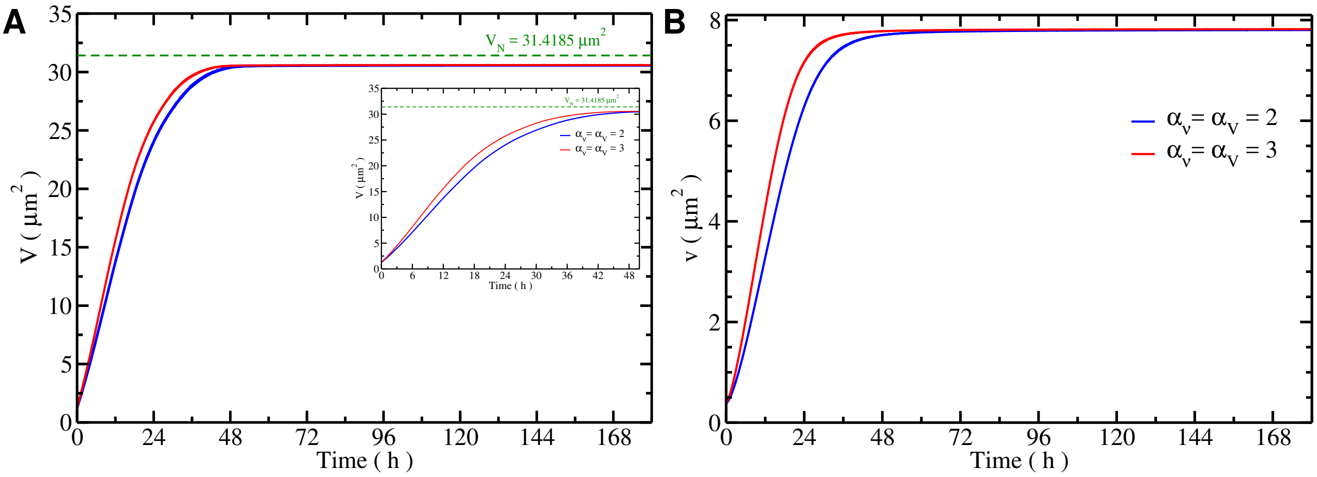


Figure 2: Temporal evolution of the overall chromosome (fig. a)) and heterochromatin (fig. b)) volumes in the cell's nucleus, without binding affinity with heterochromatin domain, for two different values of coefficient $\alpha_v$ and $\alpha_V$.

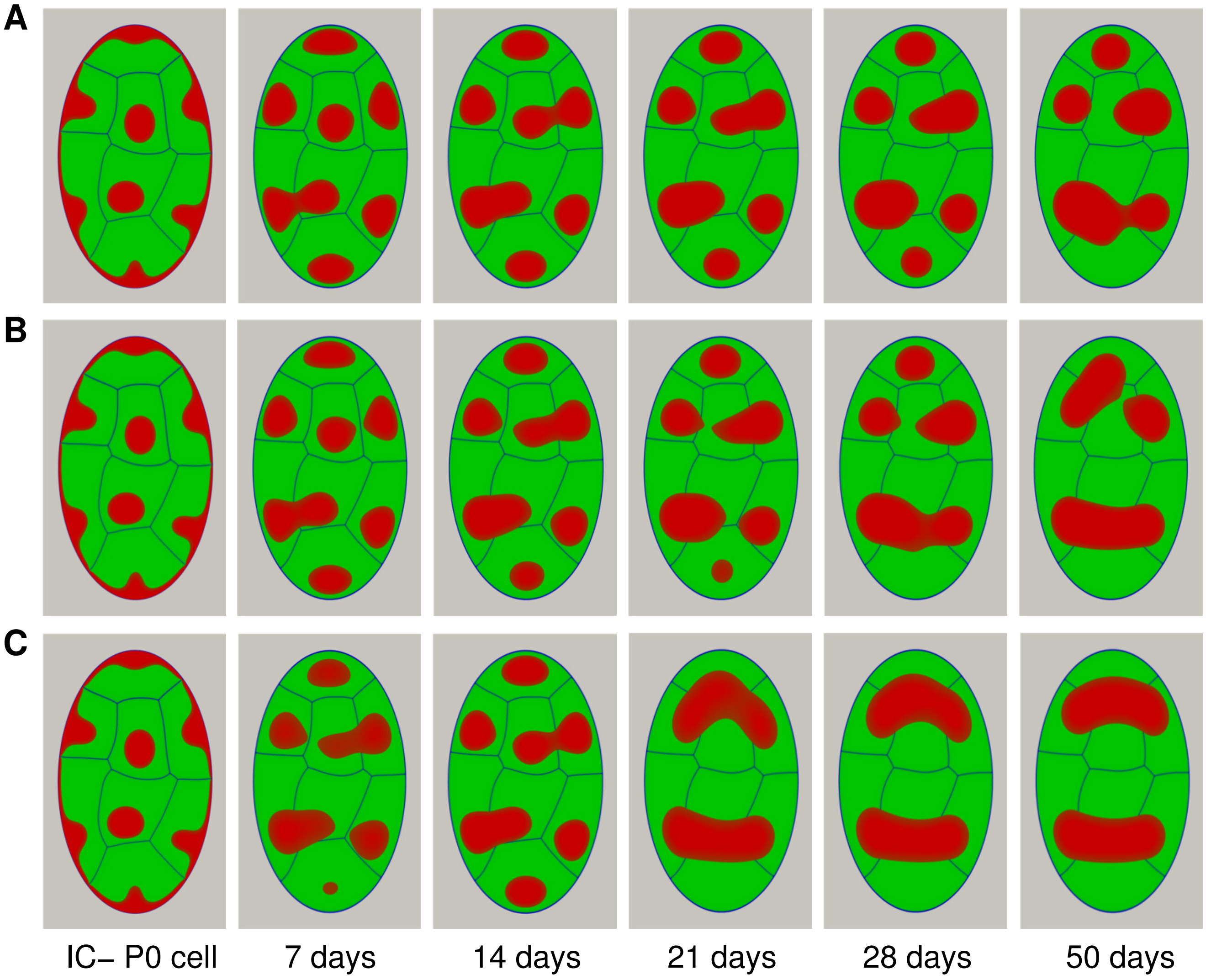


Figure 3: Effect of heterochromatin diffusion coefficient on nuclear architecture reorganization with a fixed cell's nucleus volume. The architecture obtained for a low diffusion coefficient $\epsilon_{\psi}^2 = 0.015\mu m^2$ is reported in fig a) and $0.02\mu m^2$ in fig. b), while for higher value $\epsilon_{\psi}^2 = 0.04\mu m^2$ is reported in fig. c). The heterochromatin density in cell's nucleus is fixed to 30%. The other parameters used: $\epsilon_{\varphi}^2 = 0.005\mu m^2$, $\beta_0 = 16.67$, $\beta_{\varphi} = 40$, $\beta_{\psi} = 0.1$, $\alpha_0 = 0.16$, and $\alpha_v = \alpha_V = 2$.

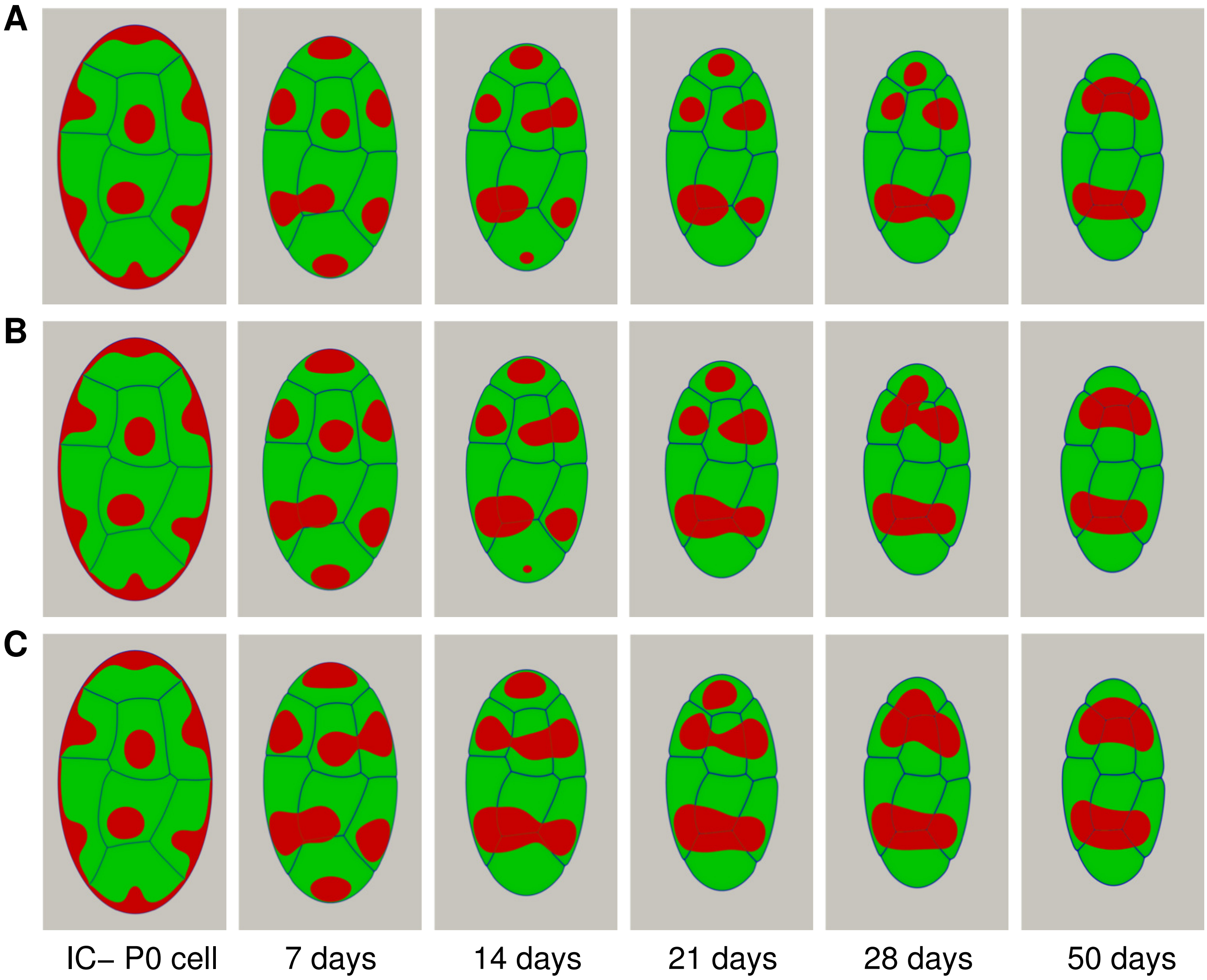


Figure 4: Effect of heterochromatin density in cell's nucleus on nuclear architecture reorganization. The architecture organization for a low density ($\rho = 25\%$) is reported in fig. a) and ($\rho = 30\%$) in fig. b) while for high value ($\rho = 35\%$) is reported in fig. c). The other parameters used: $\epsilon_\varphi^2 = 0.005$, $\epsilon_\psi^2 = 0.02$ $\beta_0 = 16.67$, $\beta_\varphi = 40$, $\beta_\psi = 0.1$, $\alpha_0 = 0.16$ and $\alpha_v = \alpha_V = 2$.

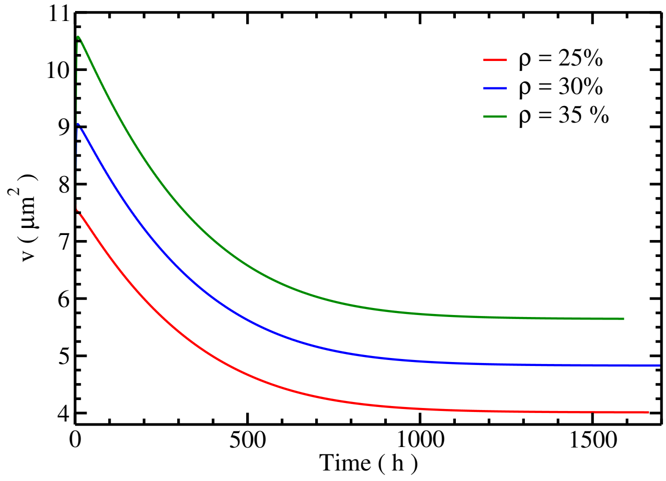


Figure 5: Temporal evolution of heterochromatin volume during the reorganization of nuclear architecture for two different value of heterochromatin density in the cell's nucleus (relaxation of heterochromatin domain: for $\rho = 25\%$ we found $a = 3.94977\mu m^2$, $b = 0.967667$ and $\tau = 305.096h^{-1}$; for $\rho = 30\%$ $a = 4.77723\mu m^2$, $b = 0.952299$ and $\tau = 303.586h^{-1}$, and we found $a = 5.53373\mu m^2$, $b = 0.957598$ and $\tau = 315.993h^{-1}$ for $\rho = 35\%$).

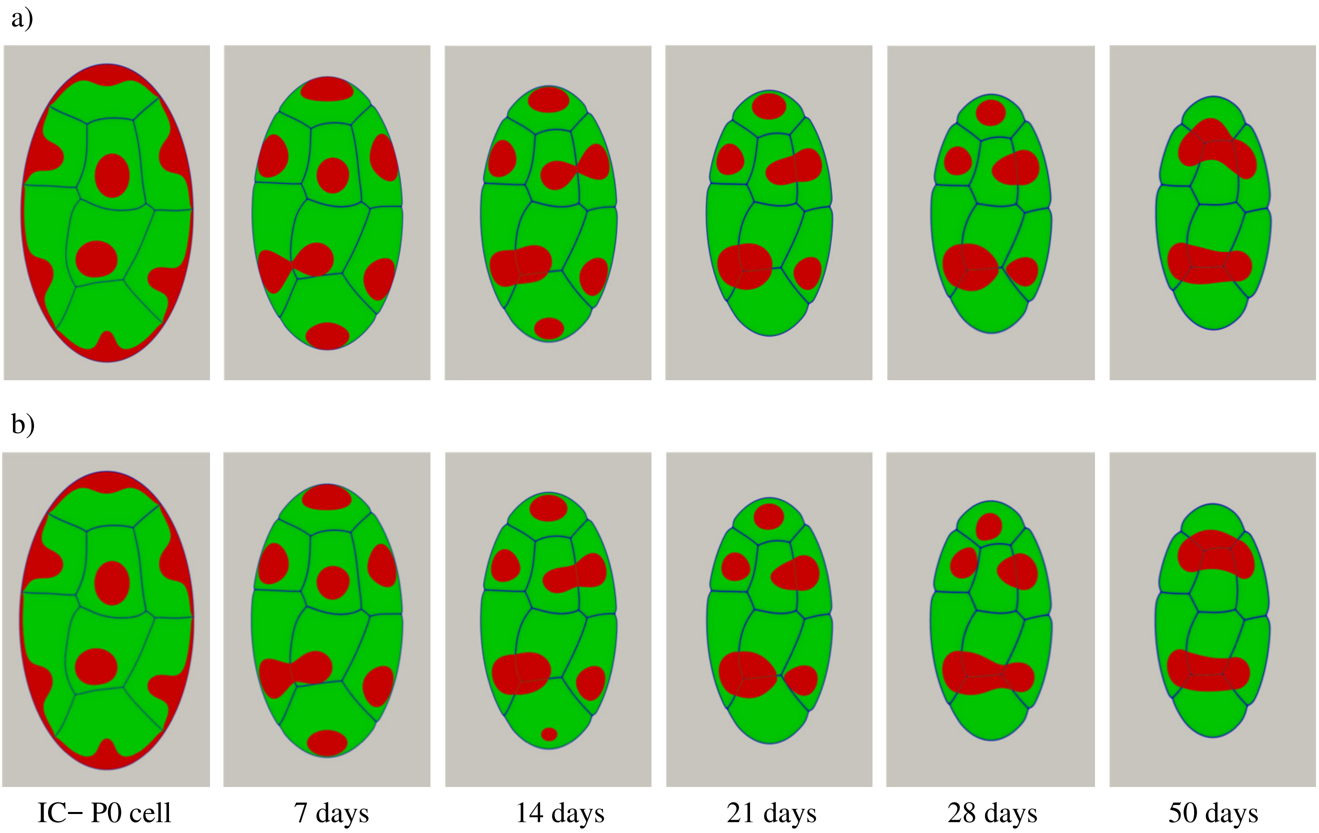


Figure 6: Effect of the heterochromatin diffusion coefficient on the reorganization of nuclear architecture accompanied by a change in the nuclear volume to reach 50% of its initial volume. The architecture organization for a low diffusion coefficient ($\epsilon_{\psi}^2 = 0.015\, \mu m^2$) is reported in fig. a) while for high value ($\epsilon_{\psi}^2 = 0.02\, \mu m^2$) is reported in fig. b). The other parameters used: $\epsilon_{\varphi}^2 = 0.005$, $\beta_0 = 16.67$, $\beta_{\varphi} = 40$, $\beta_{\psi} = 0.1$, $\alpha_0 = 0.16$, $\alpha_v = \alpha_V = 2$ and $\rho = 30\%$.

#### References


[1] A. Karma and W.-J. Rappel, "Quantitative phase-field modeling of dendritic growth in two and three dimensions," *Phys. Rev. E*, vol. 57, pp. 4323–4349, Apr 1998.

[2] U. Seifert, "Adhesion of vesicles in two dimensions," *Phys. Rev. A*, vol. 43, pp. 6803–6814, Jun 1991.

[3] J. Zhang, S. Das, and Q. Du, "A phase field model for vesicle–substrate adhesion," *Journal of Computational Physics*, vol. 228, no. 20, pp. 7837 – 7849, 2009.

[4] M. Nonomura, "Study on multicellular systems using a phase field model," *PloS one*, vol. 7, no. 4, p. e33501, 2012.

[5] S. S. Lee, S. Tashiro, A. Awazu, and R. Kobayashi, "A new application of the phase-field method for understanding the mechanisms of nuclear architecture reorganization," *J. Math. Biol.*, 2017.

[6] S. M. Allen and J. Cahn, "Microscopic theory for antiphase boundary motion and its application to antiphase domain coarsening," *Acta. Metall.*, vol. 27, p. 1084, 1979.

[7] D. Knoll and D. Keyes, "Jacobian-free newton–krylov methods: a survey of approaches and applications," *Journal of Computational Physics*, vol. 193, no. 2, pp. 357 – 397, 2004.

[8] A. E. Slaughter, J. W. Peterson, D. R. Gaston, C. J. Permann, D. Andrs, and J. M. Miller, "Continuous integration for concurrent moose framework and application development on github," *Journal of Open Research Software*, vol. 3, 11 2015.

[9] D. R. Gaston, C. J. Permann, J. W. Peterson, A. E. Slaughter, D. Andrš, Y. Wang, M. P. Short, D. M. Perez, M. R. Tonks, J. Ortensi, L. Zou, and R. C. Martineau, "Physics-based multiscale coupling for full core nuclear reactor simulation," *Annals of Nuclear Energy*, vol. 84, pp. 45–54, 2015.

[10] R. M. Tonks, D. Gaston, P. C. Millett, D. Andrs, and P. Talbot, "An object-oriented finite element framework for multiphysics phase field simulations," *Computational Materials Science*, vol. 51, pp. 20–29, 2012.

[11] B. A. Edgar and P. H. O'Farrell, "The three postblastoderm cell cycles of drosophila embryogenesis are regulated in g2 by string," *Cell*, vol. 62, no. 3, pp. 469–480, 1990.

[12] I. Solovei, M. Kreysing, C. Lanctôt, S. Kösem, L. Peichl, T. Cremer, J. Guck, and B. Joffe, "Nuclear architecture of rod photoreceptor cells adapts to vision in mammalian evolution," *Cell*, vol. 137, pp. 356–368, Apr. 2009.

[13] I. Bronstein, Y. Israel, E. Kepten, S. Mai, Y. Shav-Tal, E. Barkai, and Y. Garini, "Transient anomalous diffusion of telomeres in the nucleus of mammalian cells," *Physical review letters*, vol. 103, no. 1, p. 018102, 2009.

[14] H. Hajjoul, J. Mathon, H. Ranchon, I. Goiffon, J. Mozziconacci, B. Albert, P. Carrivain, J.-M. Victor, O. Gadal, K. Bystricky, *et al.*, "High-throughput chromatin motion tracking in living yeast reveals the flexibility of the fiber throughout the genome," *Genome research*, 2013.
